# Mass spectrometric method for the unambiguous profiling of cellular dynamic glycosylation

**DOI:** 10.1101/2020.05.04.075655

**Authors:** Asif Shajahan, Nitin T. Supekar, Han Wu, Amberlyn M. Wands, Ganapati Bhat, Aravind Kalimurthy, Masaaki Matsubara, Rene Ranzinger, Jennifer J. Kohler, Parastoo Azadi

## Abstract

Various biological processes at the cellular level are regulated by glycosylation which is a highly micro-heterogeneous post-translational modification (PTM) on proteins and lipids. The dynamic nature of glycosylation can be studied through bio-orthogonal tagging of metabolically engineered non-natural sugars into glycan epitopes. However, this approach possesses a significant drawback due to non-specific background reactions and ambiguity of non-natural sugar metabolism. Here we report a tag-free strategy for their direct detection by glycoproteomics and glycomics using mass spectrometry. The method dramatically simplifies the detection of non-natural functional group bearing monosaccharides installed through promiscuous sialic acid, GalNAc, and GlcNAc biosynthetic pathways. Multistage enrichment of glycoproteins by cellular fractionation, subsequent ZIC-HILIC based glycopeptide enrichment, and a spectral enrichment algorithm for the MS data processing enabled direct detection of non-natural monosaccharides that are incorporated at low abundance on the N/O-glycopeptides along with their natural counterparts. Our approach allowed the detection of both natural and non-natural sugar bearing glycopeptides, N and O-glycopeptides, differentiation of non-natural monosaccharide types on the glycans and also their incorporation efficiency through quantitation. Through this we could deduce some interconversion of monosaccharides during their processing through glycan salvage pathway and subsequent incorporation into glycan chains. The study of glycosylation dynamics through this method can be conducted in high throughput as few sample processing steps are involved, enabling understanding of glycosylation dynamics under various external stimuli and thereby could bolster the use of metabolic glycan engineering in glycosylation functional studies.

## Introduction

Metabolic glycan engineering (MGE), which is a technique employed to label the glycans on the living cells by treating with non-natural glycan precursors, helps in the profiling of the dynamic nature of glycosylation as it allows identification of newly synthesized glycan branches.^*1-3*^ In the process of MGE, the non-natural monosaccharides can be incorporated to the cellular glycan repertoire because of the promiscuity of glycosylation machinery. Various chemical functional groups have been successfully installed on the cellular glycoconjugates by engineering with non-natural carbohydrates via MGE.^*1*^ MGE has been exploited to label and enrich the cell surface glycome by employing analogs of the sialic acid precursor *N*-acetyl-D-mannosamine (ManNAc), *N*-acetyl-D-galactosamine (GalNAc) and *N*-acetyl-D-glucosamine (GlcNAc) in their peracetylated form.^*4-6*^ Non-natural monosaccharides incorporated into glycoproteins are detected by chemical bio-orthogonal tag-based approaches such as the Staudinger ligation and azide– alkyne cycloaddition reactions. However, these chemical probes possess limitations. In MGE, metabolic pathways allow for interconversion of unnatural sugar analogs, raising the possibility of unexpected incorporation or perturbation of cellular glycans. Therefore, it’s important to characterize the incorporation of non-natural sugars using a method which can detect both natural glycans as well as metabolically perturbed glycans simultaneously. In addition, non-specific background interactions with other reactive species in the proteome often leads to false interpretations.^*7-9*^ Several attempts to reduce these nonspecific reactions, which are proposed to be due to free radical mediated bond formation, are reported, but none of the methods were able to eliminate them completely.^*10-12*^ Even though recent strategies of using isotopic probes for chemical enrichment and isotopic recoding address the false detection of glycopeptides to a considerable extent, the use of tag-based enrichment can lead to ambiguity due to nonspecific and incomplete reactions.^*13*^

Direct tag-free mass spectrometric detection of intact glycans (glycomics) and glycopeptides (glycoproteomics) which possess these metabolically installed non-natural monosaccharides is the most viable solution for their unambiguous detection. Unambiguous and detailed mass spectrometric studies can also provide critical information on the metabolism of unnatural sugar analogs, including the positions of incorporation and possible interconversion of monosaccharides.^*14*^ However, analysis of unnatural sugar incorporation is hindered by various challenges such as the low level of incorporation of metabolically engineered sugars and preferential metabolic incorporation on a small subset of glycoforms, i.e. glycoforms other than high mannose type, in addition to other challenges associated with mass spectrometric detection of glycans and glycopeptides.^*15-17*^ Although incorporation of SiaNAz into N/O- glycans and glycosphingolipids in cells treated with Ac_4_ManNAz using tag-free mass spectrometry has been performed, detailed studies on incorporation of other frequently used azido-sugars and the peptide backbones that bear the azido-sugars are still needed for a better understanding of azido-sugar metabolism.^*18*^

We envisaged that the limitations pertaining to chemical tag-based identification of non-natural mono-saccharides on *N*/*O*- glycans could be addressed by multi-stage enrichment strategy and simultaneous reduction of the azido functional group to form an amine functional group that is more sensitive to analysis by mass spectrometry (MS) (Figures 1).^*19*^ This strategy simplifies the process of metabolically labelled glycan detection by MS through reducing the number of steps involved. It also avoids the false detection of unlabeled glycopeptides, peptides and glycans since non-specific reactions due to reporter tags do not occur.

**Figure 1:**
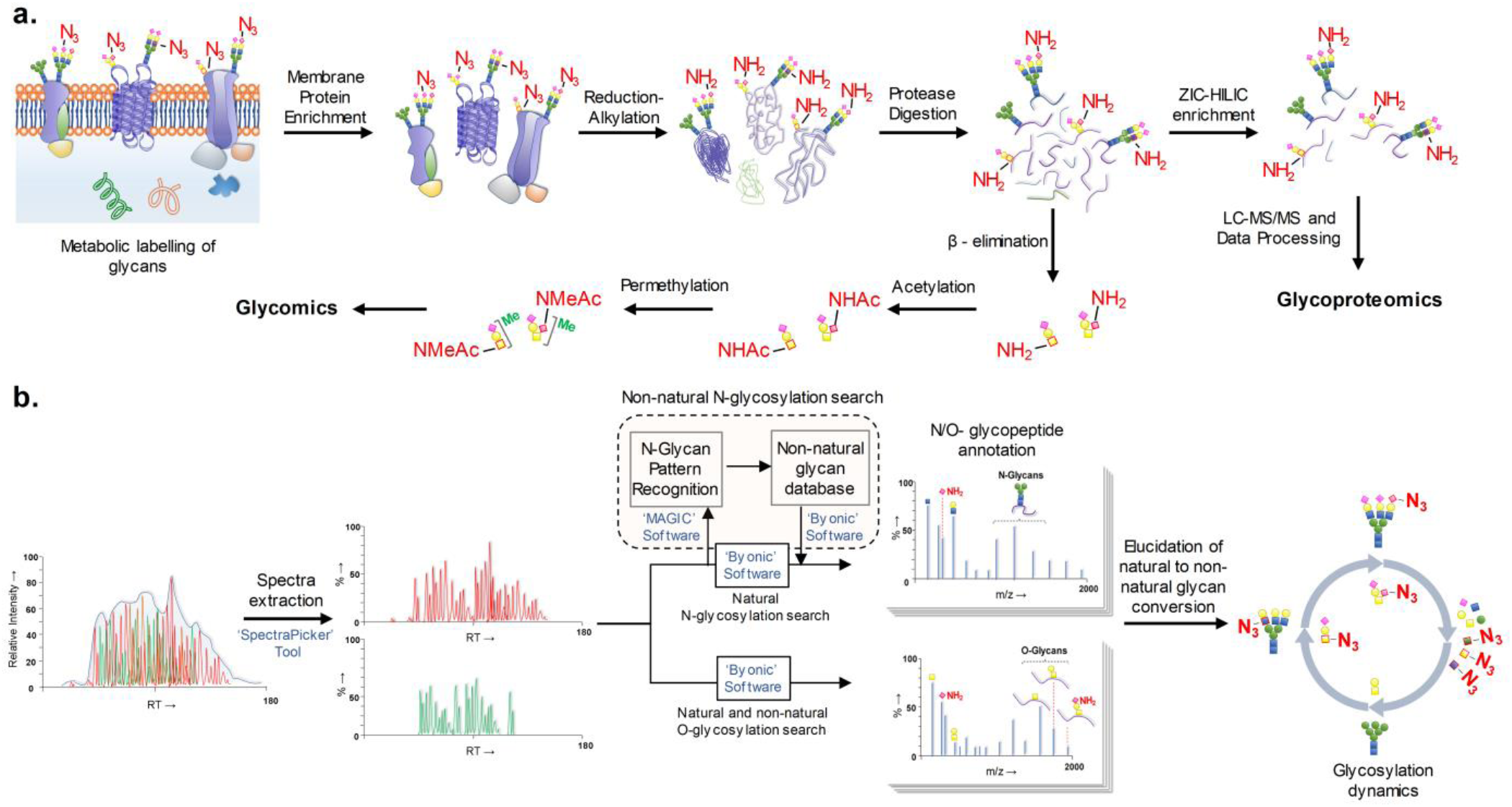
Detection of metabolically engineered non-natural glycans by chemical tag free glycoproteomics and glycomics. (a) The membrane proteins were enriched, reduced/alkylated, and digested by protease. The resulting glycopeptides were enriched for glycoproteomics by LC-MS/MS whereas the glycans were released by β-elimination followed by acetylation for glycomics; (b) For glycoproteomics, the LC-MS/MS chromatogram is extracted for the desired non-natural glycan based on the presence of oxonium ions in the spectra, detected the N-glycosylation pattern and thereby the non-natural glycan structures and searched for both natural and non-natural glycan bearing glycopeptides. O-glycopeptides are identified by the direct search of the extracted chromatogram.

## Results and Discussion

### Membrane fractionation and simultaneous enrichment of glycopeptides with and without non-natural sugars

PC-3, MCF-7, and Jurkat cells were cultured with an unnatural monosaccharide analog, either Ac_4_ManNAz, Ac_4_GalNAz or Ac_4_GlcNAz. Incorporation of the azide functional group into glycoproteins was confirmed by copper(I)-catalyzed alkyne-azide cycloaddition (CuAAC) labeling with TAMRA-alkyne followed by separation by SDS-PAGE (Figure 2a, S1).^*20, 21*^ Since *N*/*O-* glycosylation is abundant on membrane proteins, the membrane fraction was isolated from cells to enrich the metabolically engineered glycoproteins through a two-step ultracentrifugation process by differential dissolution of proteins.^*22*^ The membrane pellet was dissolved in a urea lysis buffer containing 100 mM DTT (dithiothreitol) as reducing agent, and subsequently used for the glycoproteomics and glycomics (Figure 2b, 2c, S2). The enrichment efficiency of membrane protein fractionation was evaluated by the presence of membrane localized proteins such as pan-cadherin, and cytoplasmic proteins such as GAPDH^*23*^, respectively (Figures 2d).

**Figure 2.**
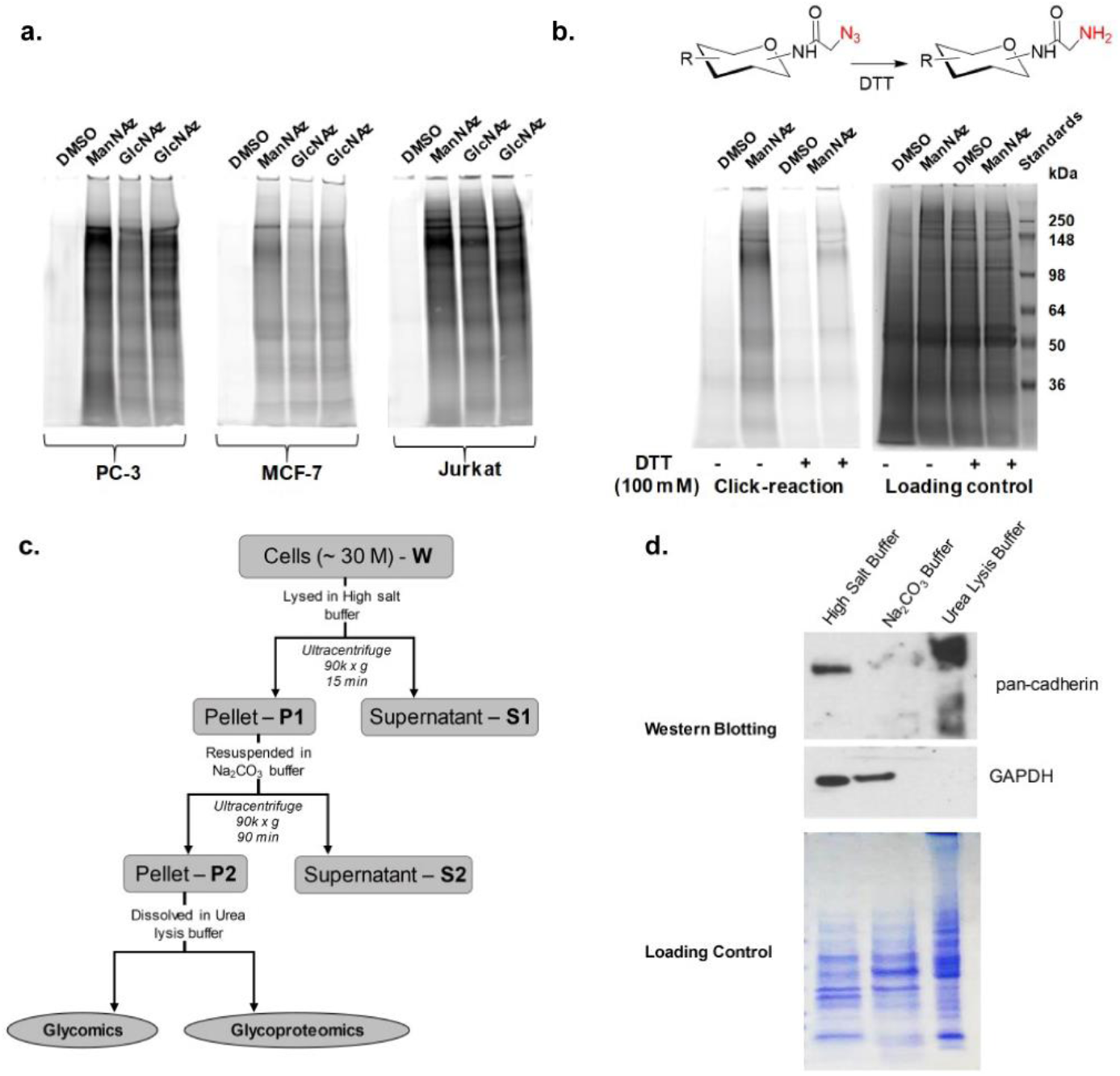
(a) Expression of azide functional group on the glycans detected by CuAAC labeling with TAMRA-alkyne (ex: 532 nm/ em: LPG/ PMT: 400); (b) the reduction of metabolically incorporated SiaNAz azide to amine in the presence of 100 mM DTT. **c**. Membrane protein fractionation strategy by lysing the cells in high salt buffer (1.0 mL; 2.0 M NaCl, 5.0 mM EDTA, pH 7.4; cytosolic-nuclear proteins removal), resuspension in sodium carbonate (Na_2_CO_3_) buffer (0.1 M Na_2_CO_3_, 1.0 mM EDTA, pH 11.3; removal of peripheral membrane proteins) and dissolving in urea lysis buffer (8.0 M urea, 1.0 M NaCl, 4 % CHAPS, 100 mM DTT, 200 mM Tris.HCl, pH 8.0); **d**. evaluation of membrane proteins fractionation by western blotting using membrane protein marker pancadherin and cytosolic protein marker GAPDH; lower panel - loading control.

The isolated proteins were proteolyzed by trypsin, enriched by a ZIC-HILIC (zwitterionic-hydrophilic interaction chromatography) enrichment and analyzed by LC-MS/MS with HCD product-triggered CID program. As expected, we detected the DTT reduction of protein disulfide bonds and simultaneous complete reduction of azide functional group to an amine (Figure S2). Also, oxonium ions of non-natural sialic acid (SiaNAz), GalNAz and GlcNAz were observed in the treated sample, but as amine modifications (*N*-aminoacetyl-neuraminic acid [Neu5NH_2_; MH^+^ = 307.1136], *N*-aminoacetyl-D-galactosamine [GalN-NH_2_; MH+ = 219.0981], and *N*-aminoacetyl-D-glucosamine [GlcN-NH_2_; MH+ = 219.0981]) on the *N*-acetyl side chain instead of the azido group (Figure 3a, 3b). Nevertheless, this does not affect the interpretation of the MS data since such modified non-natural forms of these sugars were completely absent in control (DMSO treated) samples. This observation was also confirmed when the cell lysates from MCF-7 cells were treated with DTT prior to CuAAC labeling with TAMRA-alkyne and only minimal modification with the fluorophore was observed (Figure 2b). This observation also gives some insights into the limitation of the common practice of reduction of protein disulfide by DTT while enriching the azide-modified non-natural glycans by tag-based approach as DTT-reduced amine no longer reacts with the complementary alkyne probes.^*13*^ There is a recent study in which SiaNAz incorporation into O-linked glycans was demonstrated by the direct detection of released O-glycans with SiaNAz and its partially reduced amine form Neu5NH_2_.^*18*^ Here, the O-glycans were detected in their native form by HILIC-MS and glycan structures were confirmed by matching the retention time with corresponding standard glycans. In such analysis, branching and linkage determination are not possible unless suitable standard is used. Permethylation of glycans can enable detailed mass spectrometric evaluation of branching and linkages without relying on glycan standards and can be the method of choice to determine previously unreported glycans.^*24*^ Thus, we attempted permethylation of metabolically engineered O-glycans after releasing them from the proteins by *β*-elimination. However, we could only detect natural O-glycans but not metabolically engineered one. We believed that this could be due to degradation of azide functional group during permethylation and consequent loss of it during sample processing. Considering the advantages of permethylation in detailed structural characterization of glycans, we developed a method for the simultaneous reduction, release and acetylation of O-glycans from the glycoprotein and permethylation for the glycomics profiling. Thus, the released glycans with azide functional groups were reduced to amine by DTT, selectively *N*-acetylated and detected by MS and MS^2^ after permethylation (Figures 3c, 3d, S3).

**Figure 3.**
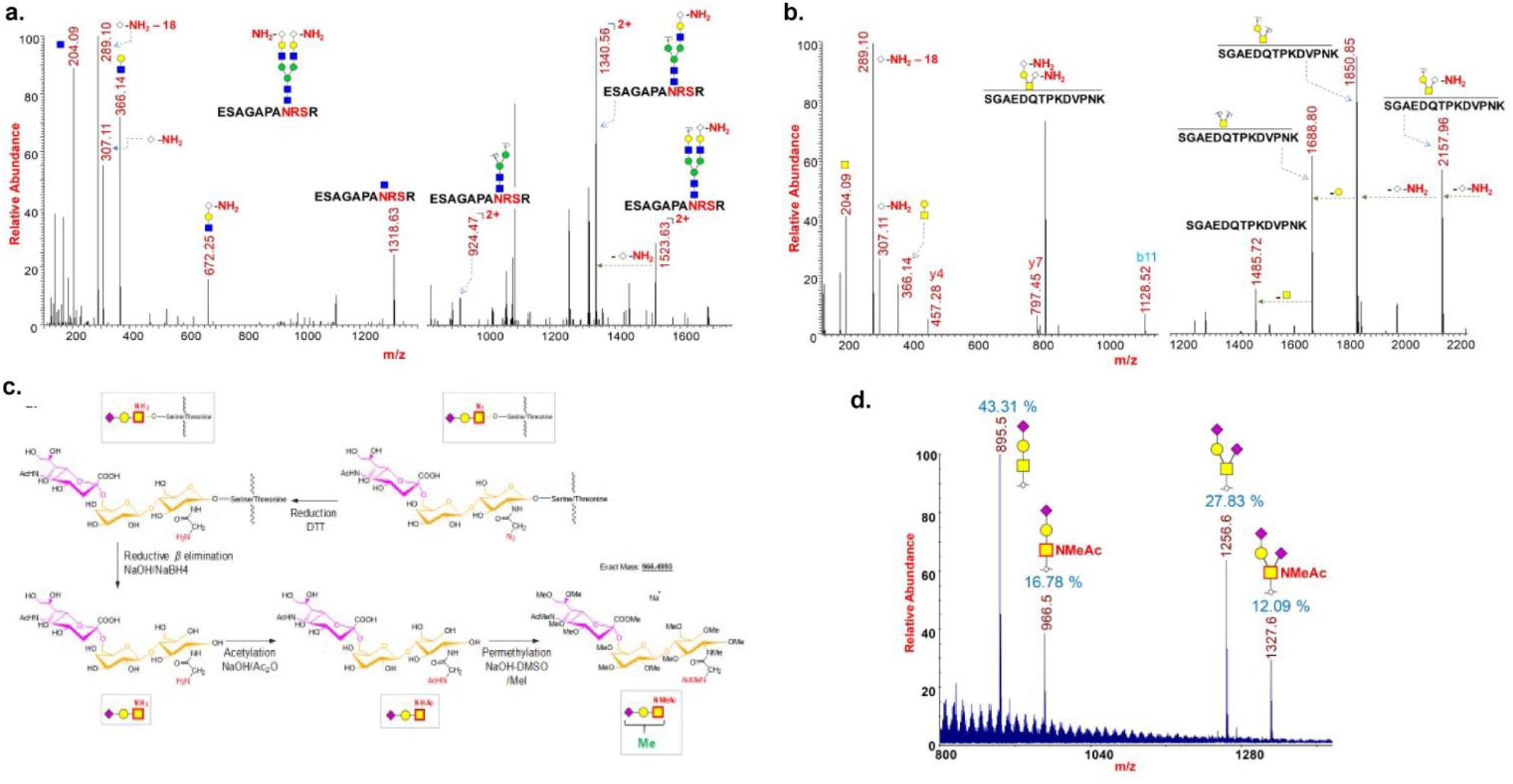
(a) Incorporation of SiaNAz on *N*-glycopeptide ‘ESAGAPANRSR’ from Neurotrophin-4 was detected in its amine form (identified by full MS, oxonium ions and neutral losses); (b) Incorporation of SiaNAz on *O*-glycopeptide ‘SGAEDQTPKDVPNK’ from Trans-Golgi network integral membrane protein 2 was detected in its amine form (identified by full MS, oxonium ions and neutral losses); (c) Scheme showing the reduction of azide on the O-glycans to amine by the treatment of DTT, reductive *β* elimination of O-glycans, acetylation and permethylation. DTT - dithiothreitol, NaOH – sodium hydroxide, NaBH_4_ - sodium borohydride, Ac_2_O – acetic anhydride, DMSO - dimethyl sulfoxide, and MeI – methyl iodide; (d) MALDI-MS spectrum of acetylated and permethylated *O*-glycans released from *O*-GalNAz treated PC-3 cells.

### Extraction of non-natural glycopeptide spectra and annotation of intact metabolically labelled N/O-glycopeptides and natural glycopeptides

While direct database search was employed for the annotation of natural glycopeptides in control and treated samples, we have developed an alternate data processing algorithm for the annotation of *N*-glycosylation with non-natural monosaccharide bearing glycans as the conventional data processing software was not effective in the glycopeptide annotation (Figure 1). We have followed an approach to filter out and categorize glycopeptide spectra unambiguously based on the presence of characteristic glycan signature ions. This enabled us to eliminate of false positive and also in reducing wrong software annotations, significantly. For this process, we first filtered the spectra bearing the signature oxonium ion for either Neu5NH_2_, GalN-NH_2_ or GlcN-NH_2_ through an in-house developed spectra extraction tool, SpectraPicker. Secondly, the spectra were searched for the ‘Peptide + HexNAc’ ion, which is the typical fragment observed in HCD MS^2^ spectra, using a *N*-glycosylation pattern recognizing software MAGIC.^*25*^ Thirdly, from the masses of ‘Peptide + HexNAc’ ion and its precursor masses, both glycan and peptide masses are deduced. Finally, a database of glycans bearing non-natural sugars were created and it is used for the search of non-natural glycan bearing *N*-glycopeptides from a list of non-natural glycan MS spectra files accurately generated based on the presence of characteristic oxonium ions by using SpectraPicker. For the annotation of *O*-glycopeptides bearing natural and non-natural sugars (Neu5NH_2_ and GalN-NH_2_), database searches were conducted on filtered MS spectra files generated by SpectraPicker based on the presence of corresponding oxonium ions.

About 629 fully characterized, manually validated sialoglycopeptides bearing SiaNAz from 256 glycoproteins were annotated on PC-3 cells treated with Ac_4_ManNAz whereas 585 Neu5Ac-containing glycopeptides from 285 glycoproteins were annotated on the DMSO-treated cells (Figure 4, S4). However, the overall number and intensity of sialoglycopeptides with SiaNAz are observed to be higher than the Neu5Ac glycopeptides in DMSO-treated cells (Figures 5a, 5b, S4). This result is consistent with the entry point of the non-natural sialic acid precursor Ac_4_ManNAz, which bypasses the master regulator of sialic acid biosynthesis, UDP-GlcNAc 2-epimerase (GNE).^*26*^ Nevertheless, about 7520 glycopeptide spectra which possess oxonium of SiaNAz and 221 spectra which possess oxonium ion of Neu5Ac were observed on PC-3 cells treated with Ac_4_ManNAz. Similarly, about 4091 glycopeptide spectra with Neu5Ac and none with SiaNAz were detected in DMSO-treated cells which pinpoints absolute elimination of false positives (Figure 4). In PC-3 cells treated with Ac_4_GlcNAz, about 387 fully characterized, manually validated sialoglycopeptides bearing Neu5Ac from 272 glycoproteins and 24 SiaNAz containing glycopeptides from 24 glycoproteins were detected. About 327 glycopeptide spectra which possess oxonium of SiaNAz and 2590 spectra which possess oxonium ion of Neu5Ac were observed on PC-3 cells treated with Ac_4_GlcNAz. (Figure 4, S5).

**Figure 4.**
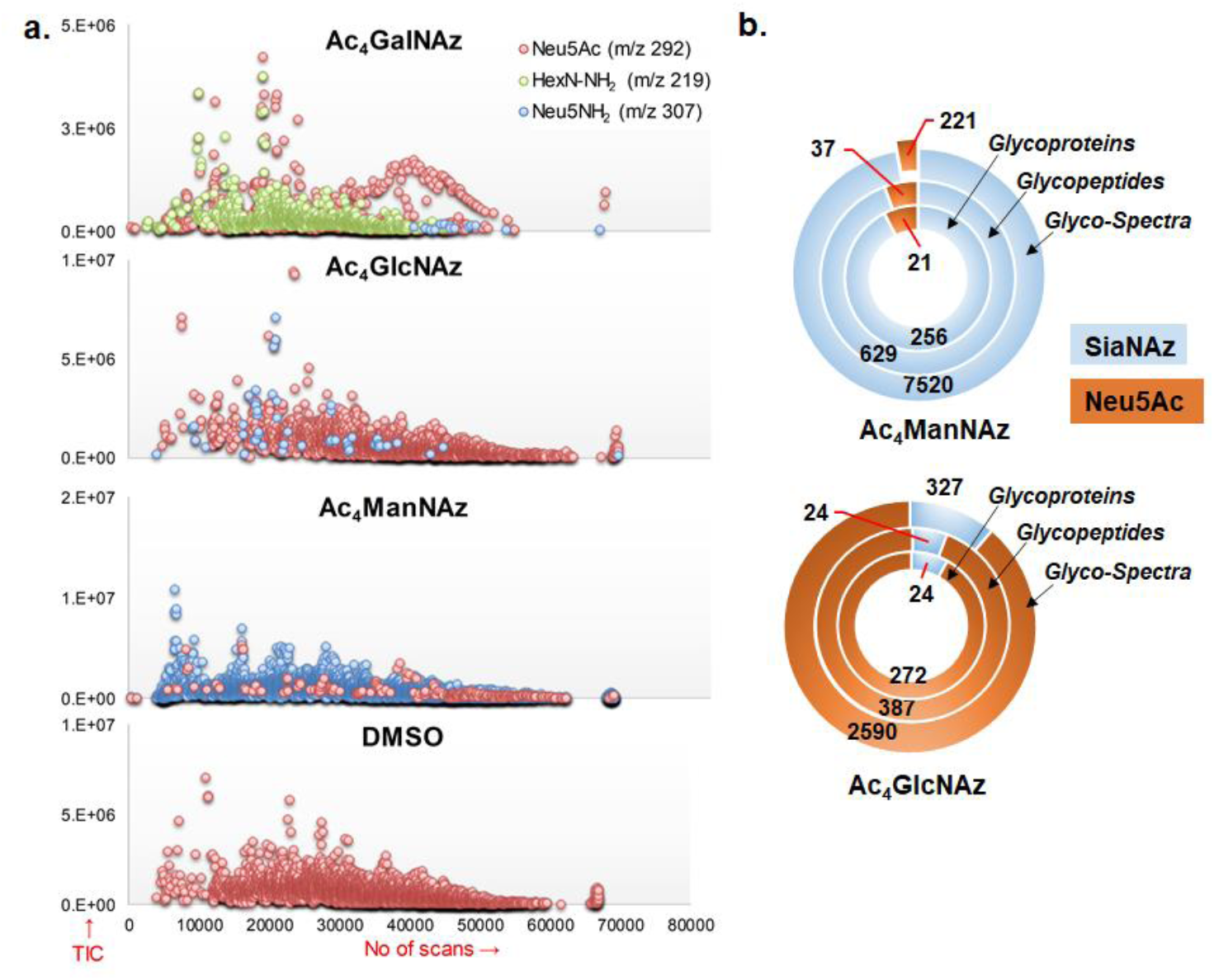
(a) Glycopeptide spectra from non-natural monosaccharide treated PC-3 cells were filtered and classified by SpectraPicker tool based on the presence of glycan oxonium ions of Neu5Ac (292), HexN-NH_2_ (219) and Neu5NH_2_ (307); Detected glycospectra, glycopeptides and glycoproteins with SiaNAz and Neu5Ac modified N-glycans. Upon Ac_4_ManNAz and Ac_4_Glc- NAc treatment on PC-3 cells.

**Figure 5.**
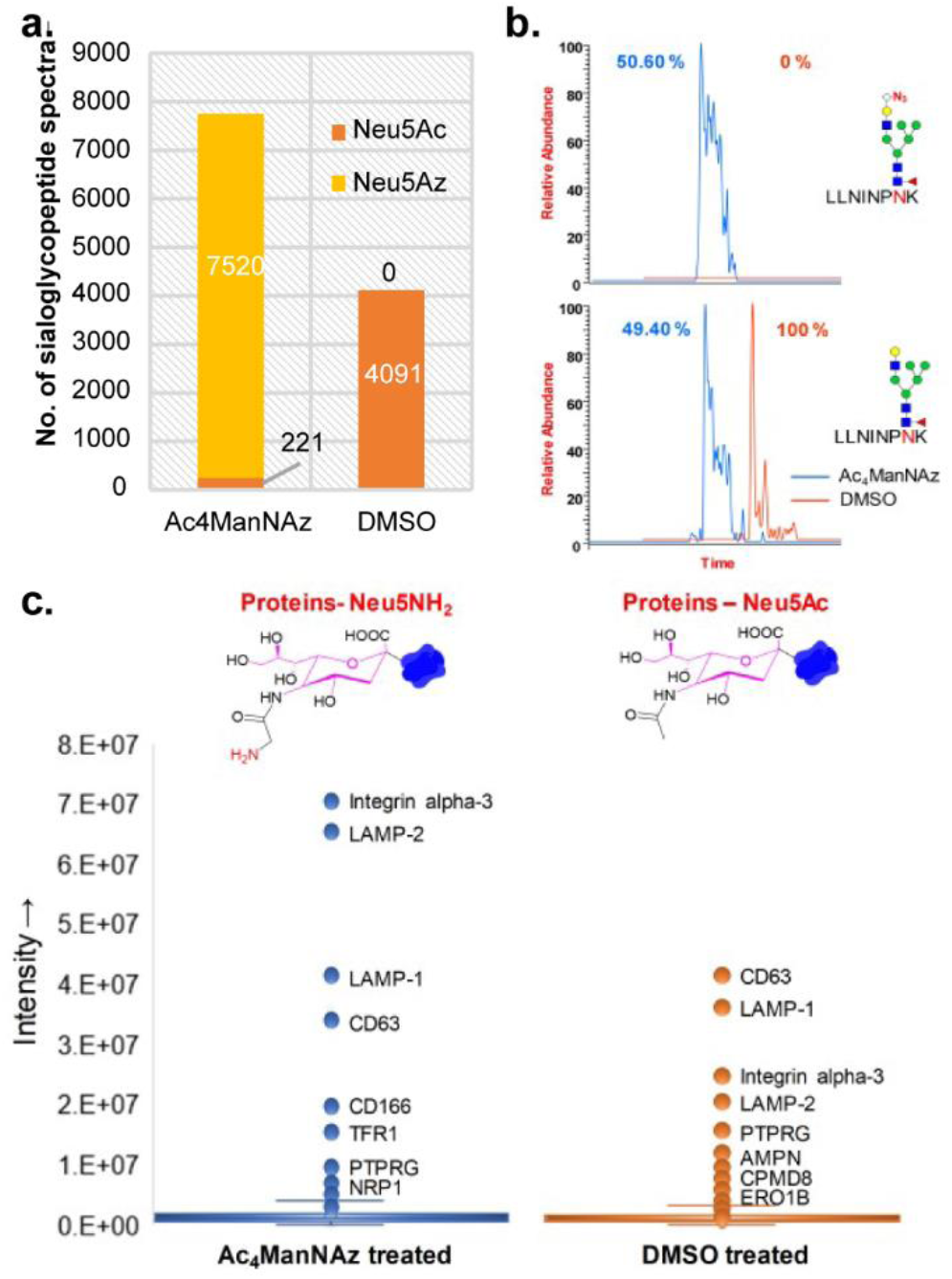
(a) No. of sialoglycopeptides (5 ppm mass accuracy and 10 % intensity cut-off) with SiaNAz and Neu5Ac detected on PC- 3 cells treated with Ac_4_ManNAz and DMSO (extracted by SpectraPicker). (b) Sialylation on the glycans on glycopeptide LLNNPNK increased with respect to vehicle upon treatment with SiaNAz; (c) Quantitative profiling of natural and non-natural sialic acid bearing sialoglycoproteins detected in Ac_4_ManNAz and DMSO treated PC-3 cells shows relatively higher level of sialoglycoproteins in Ac_4_ManNAz treated cells.

The analysis of O-glycosylation in PC-3 cells identified about 75 *O*-glycopeptides which together contain both SiaNAz and Neu5Ac upon Ac_4_ManNAz treatment. Upon Ac_4_GalNAz treatment, 103 *O*- glycopeptides which together contain both GalNAc and GalNAz were detected on PC-3 cells (Figures S6-S7) (Supp. Tables 1-6). We have quantitated the N-glycoproteins with non-natural sialic acids on Ac_4_ManNAz and natural sialic acid on DMSO treated PC-3 cells by calculating the cumulative intensity of the sialoglycopeptides. Interestingly, differences were observed on the most abundant protein for Ac_4_ManNAz treated PC-3 cells in comparison DMSO treated cells (Figure 5c). Integrin alpha-3 is observed as the most abundant non-natural sialic acid incorporated protein in Ac_4_ManNAc treated cells whereas CD63 is observed as the most abundant natural sialic acid containing protein in the DMSO control (Figure 5c). These results highlight the unique advantage of tag-less strategy as it allows simultaneous quantitation of the glycosylation dynamics within the cells as well as the biological control – comparison of natural and non-natural sugars on different samples as well as within the same sample. Moreover, a stringent comparison of each spectrum with untreated sample set is possible by evaluating the MS fragmentation spectrum by CID and HCD for signature ions and neutral losses.

In total, from the protease digest of PC-3, MCF-7, and Jurkat cells treated with non-natural monosaccharides, glycopeptides representing 49 high-mannose, complex and hybrid type N-glycan structures (13 with non-natural sugars) and 10 core-1 type sialylated *O*-linked glycans (7 with non-natural sugars) were detected (Figures 6). Each structure was assigned based on extensive manual evaluation of peptide fragments, oxonium ions and neutral losses on high resolution HCD and CID spectra (S8 to S26). Even though high-mannose type glycans were the predominant *N*-glycans observed in both treated and untreated cells, non-natural glycan modifications were not observed on them, suggesting the non-promiscuous nature of dolichol pathway.^*16, 17*^

**Figure 6:**
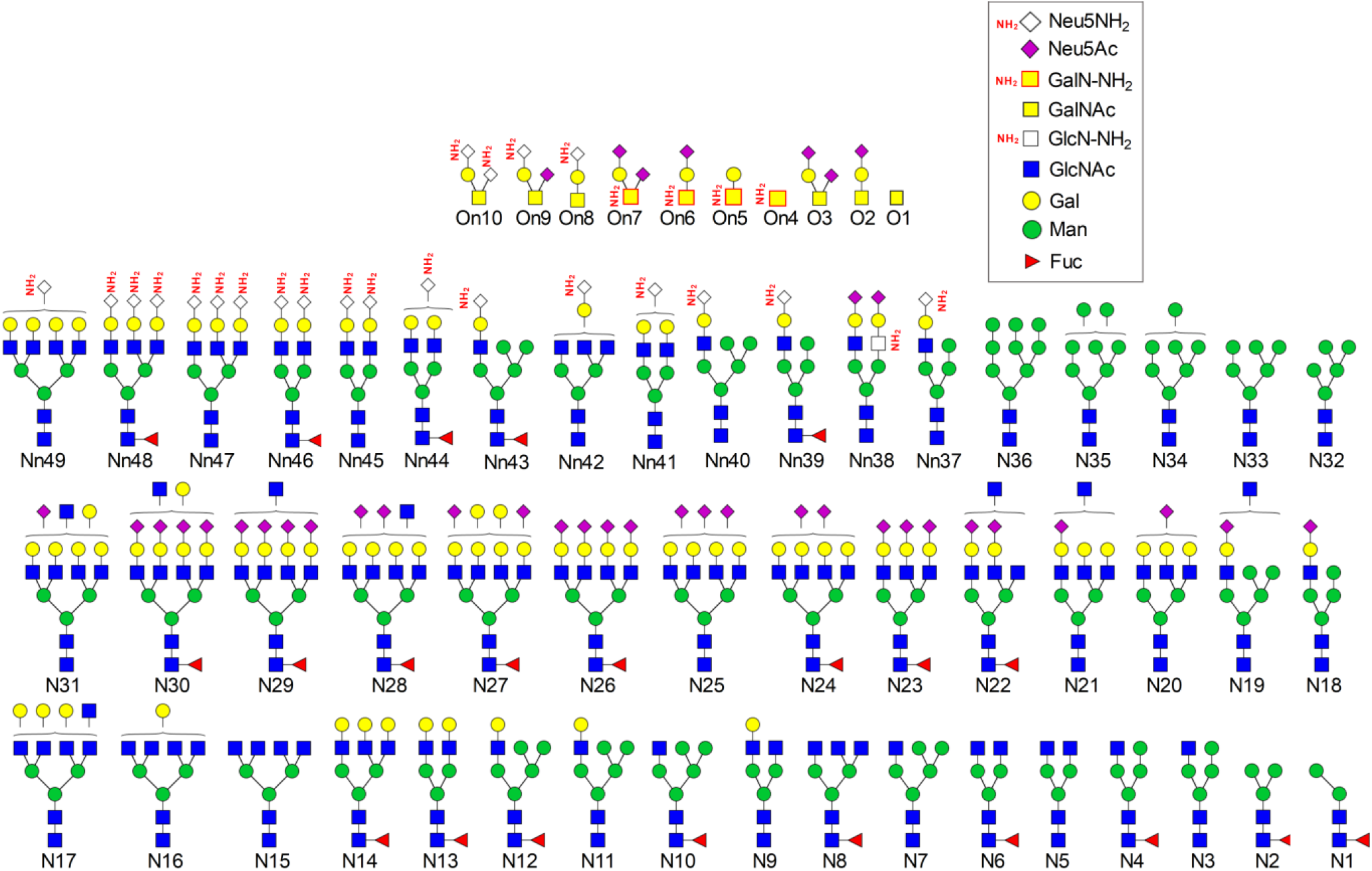
Natural (N1-36, O1-2) and non-natural (Nn37-49, On4-10) N- and O- glycans detected on PC-3 cells treated with Ac_4_ManNAz, Ac_4_GalNAz and Ac_4_GlcNAz.

### Localization of non-natural sugars on the glycan chains and validating the monosaccharide types present on glycans

In general, tag-based enrichment using biorthogonal ligation and subsequent glycopeptides bearing non-natural sugars characterization lacks information about the location of non-natural sugars on the glycan chains as the glycopeptides are released from the beads for MS characterization.^*27*^ Moreover, such tag-based detection cannot differentiate the types of monosaccharides on the glycan chains attached to glycopeptides such as sialic acid, GlcNAc or GalNAc as the glycan are released from the peptides for the characterization of glycosylation sites.^*28, 29*^ This is relevant due to possibilities of interconversion of monosaccharides during their processing through salvage pathway.^*26, 30*^ In our approach, MS based glycoproteomics was used to detect the non-natural monosaccharide directly on glycopeptides from a complex cellular proteomic mixture digest. Here, the glycan oxonium ions and neutral loss patterns on the tandem MS^n^ spectra give unambiguous proof for the presence of different types of non-natural monosaccharides and their location on glycopeptides. Thus, specific location of non-natural monosaccharides on the glycan branches (terminal or internal) and the glycan subtypes which bears the non-natural monosaccharides can be precisely identified (Figures 3a, 3b, 6). In addition to this, our approach allows multiplexing through multiple non-natural sugar treatments to the cells at the same time (Ac_4_ManNAz, Ac_4_GalNAz or Ac_4_GlcNAz) and simultaneous detection of Neu5NH_2_ GalN-NH_2_ or GlcN-NH_2_ modified glycans on the glycoproteins from the same cells (Figure 4). This is because Ac_4_ManNAz treatment leads to formation of Neu5NH_2_ oxonium ion with m/z 307.1136, GalNAz treatment leads to formation of GalN-NH_2_ oxonium ion with m/z 219.0981 only on O-glycans and GlcNAz treatment leads to formation of GlcN-NH_2_ oxonium ion with m/z 219.0981 and Neu5NH_2_ oxonium ion with m/z 307.1136 on the termini of N- and O-glycans (Figure 5a, 5c).

### Dynamics of non-natural monosaccharide incorporation through salvage pathways

Our approach of tag-free detection of metabolically incorporated sugars also revealed several interesting features of the dynamic interconversion of non-natural monosaccharides within the cellular environment (Figures 4a, 7, S4-S6). As expected, culturing PC-3 cells with Ac_4_ManNAz resulted in robust incorporation of SiaNAz into both N- and O-linked glycans, but no appreciable incorporation of GlcNAz or GalNAz was detected, as observed by others.^*18*^ In contrast, when PC-3 cells were cultured with Ac_4_GlcNAz, incorporation of GlcNAz into N-linked glycans was detected, consistent with prior observations.^*31*^ While we detected GlcNAz incorporation into the termini of N-linked glycans, we did not detect incorporation of GlcNAz to the chitobiose core. This result is consistent with prior studies of the substrate specificity of the conserved chitobiose biosynthetic pathway,^*16, 17*^ but inconsistent with a recent report that detected GlcNAz incorporation on the high mannose type of N-glycans.^*13*^ The disparity between our data and that of Woo et al. seems unlikely to be attributable to the type of N-glycans produced, since high mannose type N-glycans were predominant in all cell line we studied. Notably, PC-3 cells cultured with Ac_4_GlcNAz also incorporated SiaNAz into both N- and O-linked glycans. We confirmed the production of SiaNAz in Ac_4_GlcNAz-treated cells through DMB derivatization of sialic acids and detection by fluorescence HPLC (Supp Fig S27). Remarkably, this conversion of GlcNAz to SiaNAz and incorporation as SiaNAz was observed to be more robust than the incorporation of GlcNAz as such into *N*-glycan glycans (Figures 4, 6, 7, S5, S25, S26). Efficient metabolism of GlcNAz to SiaNAz may result from direct epimerization of GlcNAz to ManNAz by the GlcNAc-2-epimerase (RENBP) thereby bypassing GNE, which normally regulates sialic acid levels through feedback inhibition.^*26*^ This is supported by previous observations that overexpression of RENBP in Jurkat cells decreased the incorporation of SiaNAz into cell surface glycans in Ac_4_ManNAz-treated cells and that *in vitro* RENBP-catalyzed conversion of ManNAz to GlcNAz was confirmed by ^1^H-NMR.^*32*^ Finally, when cells were cultured with Ac_4_GalNAz, in addition to the expected incorporation of GalNAz into the O-linked glycans, incorporation of GlcNAz into the termini of N-linked glycans was also detected (Figures 4a, S6, S24).^*33*^ This observation is consistent with a prior report that demonstrated that UDP-GalNAz is epimerized to UDP-GlcNAz by the action of GALE, enabling incorporation of GlcNAz as the O-GlcNAc modification.^*34*^ However, while culturing cells with Ac_4_GlcNAz led to SiaNAz incorporation into both N- and O-linked glycans, no SiaNAz was detected when cells were cultured with Ac_4_GalNAz. One possible explanation for this observation is that conversion of UDP-GlcNAz to ManNAz is inefficient due to both feedback inhibition of and substrate specificity of GNE. Alternately, it may be that only a relatively small fraction of added Ac_4_GalNAz is converted to UDP-GlcNAz, thereby limiting flux to SiaNAz. Taken together, the data presented here provide a comprehensive picture of azido-sugar metabolic outcomes.

**Figure 7:**
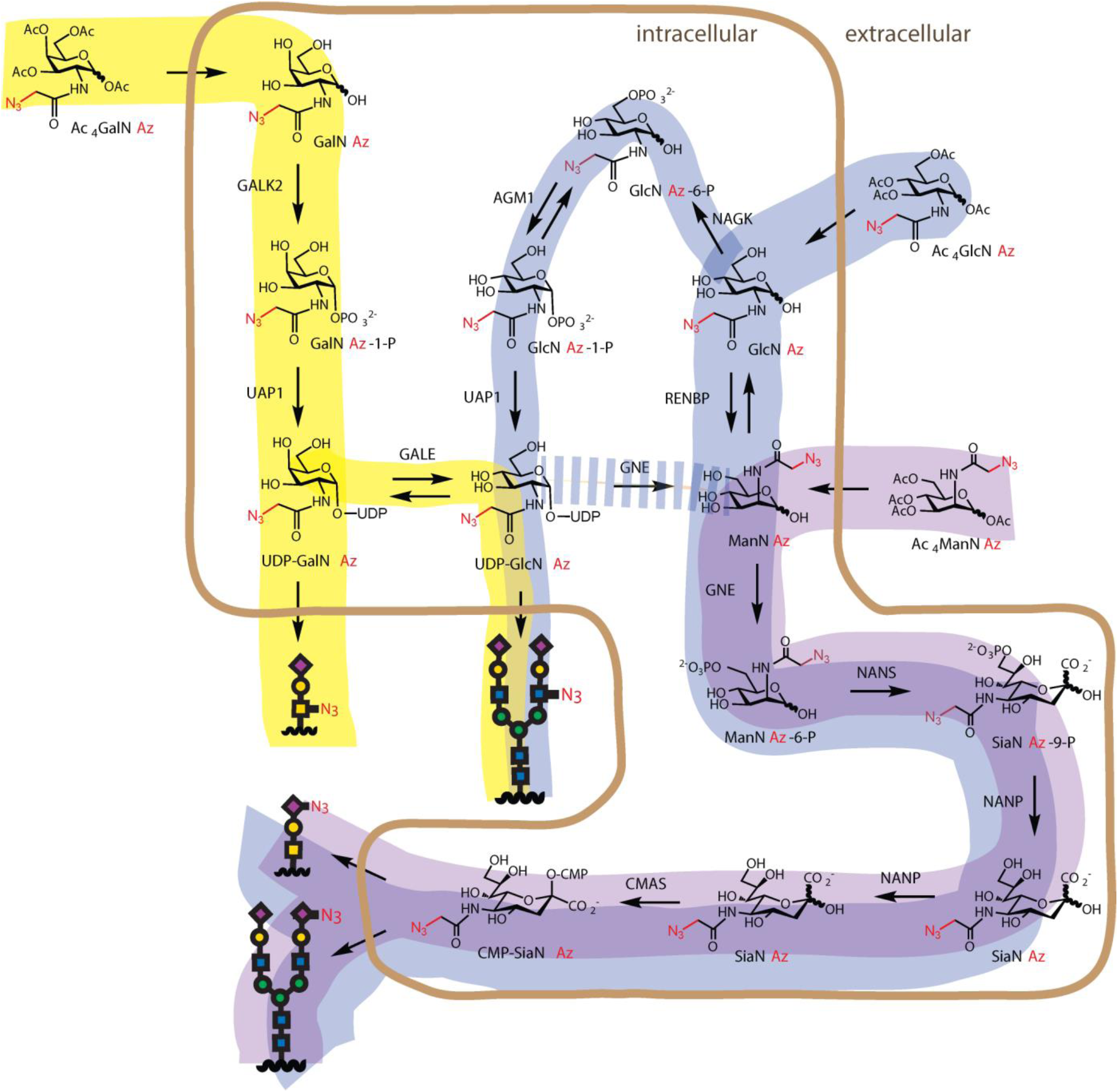
Dynamic conversion of non-natural monosaccharides within the cellular environment and the non-natural metabolite utilization pathway observed in PC-3, MCF-7 and Jurkat cells. *In vivo* - incorporation of Ac_4_ManNAz as SiaNAz, Ac_4_GalNAz as GalNAz and GlcNAz (epimerization of UDP-GalNAz by GALE enzyme), and Ac_4_GlcNAz as GlcNAz and SiaNAz (facile conversion of GlcNAc to ManNAz).

This information may be especially useful in the interpretation of experiments conducted using these important tools, as metabolic engineering with azido sugars has been employed in many studies with various applications, such as the dynamics of O-GlcNAcylation and *in vivo* imaging of glycosylation events.^*3, 9*^ However, despite its straightforward concept, the metabolism of these azido sugars has the potential to pose pitfalls for studies using these tools. Many important questions still need to be answered to better understand and interpret results from MGE, such as whether the non-natural sugars undergo interconversion just like natural monosaccharides, how the change in flux of sugar donors and the different tolerance of glycan-processing enzymes for non-natural sugars will shape the cellular glycome and into which exact positions these non-natural sugars are incorporated.^*1, 14*^ Using PC-3 cells as an example, our data answered part of the questions for azido sugars and sheds light on the “black box” of MGE. It’s important to note, however, that most of observations reported here are from a single cell line and may not all be directly transferable into other cell lines. Nevertheless, our tag-free approach provides an opportunity to look deeper into the dynamics of azido sugar incorporation and may be used to characterize the possible interconversion and incorporation of other non-natural sugars and in different cell lines.

### Significance of studying the dynamic glycosylation processes

Interestingly, when the quantitative sialoglycoproteomics data from PC-3 cells (Figure 5c) was evaluated for their involvement in diseases, we found pathway enrichment of Pi3K and FGR3 on only Ac_4_ManNAz and not on DMSO-treated case which implicates the role of sialylation dynamics on protein associated with these pathways which are known to promote prostate cancer (Figure S28). Thus, the increased sialylation induced by delivery of a protected ManNAc analog (Ac_4_ManNAz in this case), may mimic the hypersialylation state often observed in prostate cancer, as well as other cancers.^*35, 36*^ Direct MS detection of intact glycopeptides with non-natural sugars is the only method which can be employed for the quantitative determination of non-natural sugars with any functional groups. Since the technical bias due to chemical tags can be eliminated by tag-free method and the method involves facile sample processing, such strategy coupled with development of deep learning-based algorithms can potentially automate the study of complex roles glycosylation dynamics plays in cellular physiology.

## Methods

### Enrichment of membrane proteins from the cells and protease digestion

Membrane proteins were enriched from the cells before proceeding with protease digestion and glycopeptide enrichment. The cells (30 × 10^6^) were pelleted (W, whole cell lysate was prepared from an aliquot) (1000 x g for 5 min) and homogenized in high salt buffer (1.0 mL; 2.0 M NaCl, 5.0 mM EDTA, pH 7.4) by passing through 26-gauge syringe for 10 times. The homogenate was probe sonicated for 5 min using 10 sec intervals pulsed program and ultracentrifuged at 90000 X g for 15 min at 4 °C. The supernatant (S1) was separated and the pellet (P1) was re-suspended in 1.0 mL sodium carbonate buffer (0.1 M Na_2_CO_3_, 1.0 mM EDTA, pH 11.3) and kept for 30 min on ice. Further, the suspension was ultracentrifuged at 90000 x g for 90 min at 4 °C. The supernatant (S2) and the pellet (P2) was separated and the pellet containing membrane fraction was dissolved in 1.0 mL urea lysis buffer (8.0 M urea, 1.0 M NaCl, 4 % CHAPS, 100 mM DTT, 200 mM Tris.HCl, pH 8.0) and heated at 50 °C for 45 min for denaturation. After denaturation 55 mg of IAA was added to the samples and incubated at room temperature in dark for 45 min. The proteins were precipitated by sequential addition of chloroform (1.5 mL), methanol (4.5 mL) and water (5.0 mL). The precipitated protein was separated by centrifugation at 3000 X g for 5 min, washed once with 500 µL of methanol, centrifuged, and resuspended in 50 mM ammonium bicarbonate buffer (pH 8.0).The proteins were digested by adding 25 µg each of trypsin-LysC protease cocktail and incubated at 37 °C for 24 h. MS compatible protease inhibitor cocktail was added to all buffers just prior to the addition.

### Enrichment of glycopeptides from the protease digest

The glycopeptides were enriched by ZIC-HILIC method using ProteoExtract® Glycopeptide Enrichment Kit (Millipore, 72103-3) with a modified procedure. The concentration of peptides in the protease digest was estimated by nanodrop spectrophotometer and 100 µg of peptides in 30 µL were mixed with 150 µL of binding buffer.150 µL of ZIC-HILIC resin was centrifuged to remove the solvents, the sample was added to it and vortexed for 20 min at room temperature. The samples with resin was centrifuged, supernatant was discarded, the resin was resuspended in 450 µL washing buffer and incubated at room temperature for 10 min. The wash buffer was removed by centrifugation and the washing procedure was repeated two more times. Subsequently, the enriched glycopeptides were eluted by adding 225 µL of elution buffer followed by 225 µL of 0.1 % formic acid, incubating for 5 min at room temperature in each case. The eluted fractions were combined and evaporated to dryness by speed vacuum. The enriched glycopeptides were dissolved in 25 µL of 0.1 % formic acid for the subsequent LC-MS/MS analysis.

### Data acquisition of protein digest samples using nano-LC-MS/MS

The glycopeptides were analyzed on an Orbitrap Fusion Tribrid mass spectrometer with HCD, CID and ETD fragmentation option, connected to a Dionex Ultimate 3000 LC-MS/MS system. Acclaim PepMap® nano-LC columns (Thermo Scientific; Cat No. 164568) of 150 mm length with 75 µm internal diameter (id), filled with 3 µm, 100ଊ C18 material (reverse phase) were used for chromatographic separation of samples. Solvent A consisted of 0.1% formic acid (Fluka) in LC-MS grade water (Sigma Aldrich), and solvent B consisted of 0.1% formic acid, 80 % acetonitrile (Sigma Aldrich) and 20 % LC-MS grade water. Samples were run for 180 min at 300 nL/min with gradients as follows, 180 min LC method: 5-30 % solvent B within 90 min; 30-60 % solvent B within 60 min; 60-99 % solvent B within 15 min; 99-80 % solvent B within 5 min; 80-10 % solvent B within 5 min and 10-5 % solvent B within 5 min. All instrument methods for the mass spectrometer were set up in data dependent acquisition mode with an automatic gain control (AGC) target value of 5 Χ 10^5^. After the precursor ion scan at 120,000 resolution in Orbitrap analyzer, intense precursors were selected for subsequent fragmentation using HCD or CID (product-triggered based on the presence of glycan oxonium ions in the HCD) within 3 sec at a normalized collision energy of 28 and 35, respectively. For internal mass calibration 445.120025 ion was used as lock mass with a target lock mass abundance of 0 %. Charge state screening was enabled, and precursors with unknown charge state or a charge state of +1 were excluded. Dynamic exclusion was enabled (exclusion size list 100, exclusion duration 30 s). The fragment ions were analyzed in the Orbitrap at 15000 resolution.

### Data analysis for quantitative glycoproteomics through glycopeptide spectra enrichment (using SpectraPicker tool)

The identification of the LC-MS/MS glycopeptide spectra file was performed using the commercial software Byonic™ 2.3 (https://www.proteinmetrics.com/products/byonic/). Since Byonic™ with its integrated glycan database does not support the modifications on the glycan chains, the identification of the non-natural glycans does not provide good result. We developed a workflow that is based on the separation of the natural and non-natural glycan spectra and separate annotation. To this end, we developed software called SpectraPicker (freely available at our source code repository https://github.com/ReneRanzinger/SpectraFiltering/) that allows extracting MS scans based on user defined key ions in the MS^2^ data. To extract the scans for our tagged non-natural glycans all scans containing the oxonium ions (m/z 307.1136 for Neu5NH_2_ and 219.0981 for HexN-NH_2_ +/- 5ppm accuracy) with an intensity value of more than 10 % of the highest MS^2^ intensity have been extracted and saved into a separate filtered data file. In the extracted MS^2^ scans, the fragments ‘Peptide + HexNAc’ on the HCD fragmentation spectra were detected using a pattern recognizing software, MAGIC. Based on these fragments and the precursor m/z this software allows the calculation of the non-natural glycan masses and the generation of a new glycan database based on these masses. Subsequently, the newly generated non-natural glycan database is used in Byonic™ for the annotation of the non-natural monosaccharide bearing glycoproteins found in the separated spectra file. In Byonic™, the UniProtKB human reviewed *N*-glycoproteome proteome data set (download date 14.10.2018) was used in combination with the generated glycan mass database. Oxidation of methionine and carbamidomethylation of cysteine was also used as variable modifications. A precursor ion tolerance of 5 ppm and fragment ion tolerance of 15 ppm was set for the search with up to two missed cleavage for the target enzyme trypsin. The natural *N*-glycan bearing glycoproteins in the treated and control (DMSO) cells were detected by direct search of the filtered spectral files (based on oxonium ions m/z 204.0865 for HexNAc) using Byonic™ software. For the glycoprotein quantification (only sialoglycoproteins), the scan number of manually verified annotated *N*-glycopeptides were matched with the ion intensity of the corresponding spectra and subsequently the cumulative intensity of each protein was calculated from the ion intensity of glycopeptides.

For the annotation of natural and non-natural monosaccharide bearing *O*-glycoproteins, the search of spectra file extracted (SpectraPicker; 5 ppm accuracy; 10 % intensity threshold) based on the presence of corresponding oxonium ions (m/z 307.1136 for Neu5NH_2_ and 219.0981 for HexN-NH_2_) was conducted using Byonic 2.3 software against UniProtKB human reviewed proteome data set (download date 14.10.2018). Common core-1 type *O*-glycans with and without non-natural sugar modifications, oxidation of methionine and carbamidomethylation of cysteine were used as variable modifications. A precursor ion tolerance of 5 ppm and fragment ion tolerance of 10 ppm was set for the search with up to two missed cleavage for the target enzyme trypsin.

### Release of O-glycans by β-elimination and analysis of glycans by mass spectrometry after derivatization by permethylation (Glycomics)

The proteins were precipitated from the urea lysis buffer, washed with methanol and suspended in 50 mM sodium hydroxide (250 µL) (Fisher Scientific; Cat. No. 3727-01). Sodium borohydride (19 mg) (Acros organics; Cat. No. 200050250) dissolved in 50 mM sodium hydroxide (250 µL) was added to the sample and incubated at 45 C for 18h. After the incubation, the sample was cooled in an ice bath and 50 µL acetic anhydride (Sigma Aldrich; Cat. No. 320102) was added. After 10 min incubation one more crop of acetic acid was added to the sample and incubated in ice for 50 more minutes. The sample containing released, acetylated O-glycans were passed through H^+^ acid foam (Biorad; Cat. No. 142-1441) and lyophilized. The borates were removed by the addition of methanol: acetic acid mixture (9:1) and passing a stream of Nitrogen gas. After the removal of borates, the O-glycans were permethylated by the addition of DMSO (200 µL), NaOH-DMSO base (300 µL), and methyl iodide (100 µL) and incubating for 10 min. The permethylated glycans were extracted by dichloromethane, dried and analyzed by MALDI-MS and ESI- MS^n^ after re-dissolving in methanol.

## ASSOCIATED CONTENT

## Supporting Information

The Supporting Information is available free of charge on the ACS Publications website. Experimental methods, characterization and additional results (PDF and .xlsx)

SpectraPicker software tool is freely available at our source code repository https://github.com/ReneRanzinger/SpectraFiltering/

The raw data files can be accessed from glycopost repository - https://glycopost.glycosmos.org/preview/15840403535d8144a176e02; Pin: 6599

## AUTHOR INFORMATION

### Author Contributions

A.S. and P.A. conceived of the paper; A.S., N.S., H.W., A. W. and G.B. performed experiments; A.K., M.M. and R.R. developed the software tool, A.S., N.S., H.W. and J.K. analyzed the data and wrote the paper; P.A. and J.K. monitored the project.

### Notes

The authors declare no competing financial interests.

## ACKNOWLEDGMENT

We thank Prof. Russell W. Carlson, UGA for the thorough review of the manuscript and the members of analytical services and training lab, CCRC (Complex Carbohydrate Research Center) for helpful discussions and suggestions. This research was supported by the National Institutes of Health (NIH) funded research grants S10OD018530 to P.A. at CCRC, R21DK112733 to J.K. and McKnight Fellowship (Department of Biochemistry, UT Southwestern Medical Center) to H.W..

## TOC

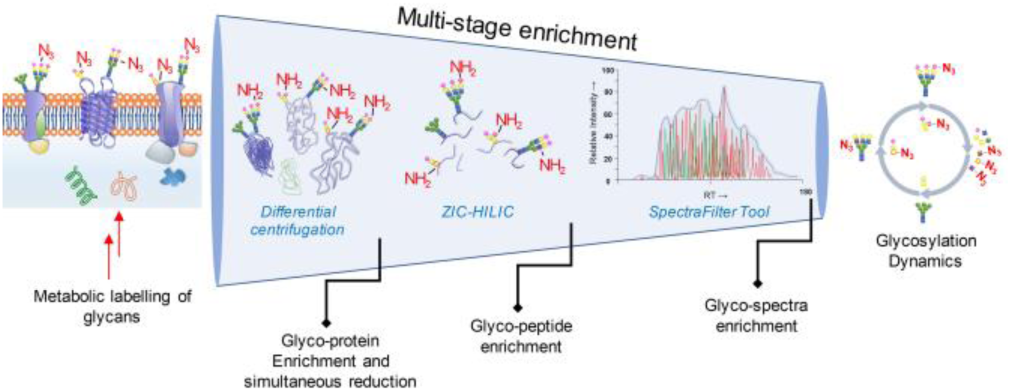

