## Supplementary information for "Mass spectrometric method for the unambiguous profiling of cellular dynamic glycosylation"


| Contents | Pages or location |
|---|---|
| Supp Methods | Pages 1 to 4 |
| Supp Figures 1 to 28 | Pages 5 to 23 |
| Supp References | Page 23 |
| Supp Tables (in .xlsx) | |
| Table 1 | Table1_Ac4ManNAz_PC3_N-Glycopeptides_NoSiaNAz_Byonic-Manual_16Sep2019 |
| Table 2 | Table2_Ac4ManNAz_PC3_N-Glycopeptides_SiaNAz_Byonic-Manual_16Sep2019 |
| Table 3 | Table3_Ac4ManNAz_PC3_O-Glycopeptides_Byonic-Manual_16Sep2019 |
| Table 4 | Table4_Ac4GalNAz_PC3_O-Glycopeptides_Byonic-Manual_16Sep2019 |
| Table 5 | Table5_DMSO_PC3_N-Glycopeptides_NoSiaNAc_Byonic-Manual_16Sep2019 |
| Table 6 | Table6_DMSO_PC3_N-Glycopeptides_SiaNAc_Byonic-Manual_16Sep2019 |

### Material and methods

#### Materials and Chemicals

Dimethyl sulfoxide (DMSO) was purchased from Fischer Scientific (Cat. No. BP231-100). $Ac_4$ManNAz (Cat. No. 1084-25 and 1084-100) and $Ac_4$GlcNAz (Cat. No. 1085-25 and 1085-100) were both purchased from Click Chemistry Tools. $Ac_4$GalNAz was synthesized by Kumar Paulvannan from Tandem Sciences, Inc. or purchased from Click Chemistry Tools (Cat. No. 1086-25). Stock concentration of $Ac_4$ManNAz, $Ac_4$GlcNAz and $Ac_4$GalNAz is 500 mM in DMSO. TAMRA-alkyne was purchased from Click Chemistry Tools (Cat. No. TA108-5), and stock concentration is 10 mM in DMSO. Tris[(1-benzyl-1-H-1,2,3-triazol-4-yl)methyl]amine (TBTA) (Cat. No. 678937-50MG), tris[(2-carboxyethyl)phosphine hydrochloride (TCEP) (Cat. No. C4706-2G) and copper(II) sulfate pentahydrate (Cat. No. C7631-250G) were purchased from Sigma-Aldrich. The stock concentration of TBTA, TCEP and copper(II) sulfate pentahydrate is 10 mM in DMSO, 50 mM in water and 50 mM in water, respectively. TCEP and copper(II) sulfate pentahydrate stocks are always made in fresh right before the CuAAC labeling. Methanol (Cat. No. A452-4), acetonitrile (Cat. No. A998-4), chloroform (Cat. No. C298-500) and dithiothreitol (DTT) (Cat. No. BP172-25) were purchased from Fisher Scientific. Stock concentration of DTT is 1 M in water. All stocks are stored at -20 °C unless otherwise specified. Urea (Cat. No. U0631), dithiothreitol (DTT Cat. No. 43815), iodoacetamide (IAA Cat. No. I1149), CHAPS (Cat. No. C3023), EDTA (Cat. No. ED4SS) and MS compatible protease inhibitor (Cat. No. MSSAFE) were purchased from Sigma-Aldrich. $Na_2CO_3$ (Cat. No. 3604-01) was purchased from J. T. Baker. Sequencing-grade modified trypsin-LysC (Cat. No. V1891) was purchased from Promega. Antibodies were obtained from Santa Cruz Biotechnology, TX (GAPDH, Cat No. sc-47724) and Cell Signaling, MA (Pan-cadherin; Cat No. 4068). 4,5-methylenedioxy-1,2-phenylenediamine dihydrochloride (DMB) was purchased from Sigma (Cat. No. 66807-50MG) and added freshly within an hour before the reaction. All other chemical reagents were purchased from Sigma-Aldrich, unless otherwise mentioned. Fisher Scientific 550 Sonic dismembrator was used for probe sonication. Ultracentrifugation was performed on a Beckmann Optima L-90K ultracentrifuge using 50.4 Ti rotor. The protein and peptide concentration were estimated using a Thermo nanodrop 2000c spectrophotometer. Mass spectrometric data acquisition was performed in the positive ion mode on a Thermo Scientific LTQ Orbitrap Fusion Tribrid mass spectrometer coupled with Dionex Ultimate 3000 LC system. Data analysis was performed using Byonic 2.3 software and by manual method using Xcalibur 3.0.

#### Cell lines and cell culture reagents

Jurkat cells were obtained from Dr. Kim Orth, UT Southwestern Medical Center. MCF-7 and PC-3 cells were purchased from ATCC. Jurkat cells were cultured in RPMI 1640 medium (Thermo Fisher Scientific, Cat. No. 11875093) containing 10% fetal bovine serum (Thermo Fisher Scientific, Cat. No. 16000044), 10 mM HEPES (Thermo Fisher Scientific, Cat. No. 15630080), 100 μg/mL streptomycin and 100 units/mL penicillin (Thermo Fisher Scientific, Cat. No. 15140122). MCF-7 cells were cultured in high glucose DMEM medium (Thermo Fisher Scientific, Cat. No. 11965092) containing 10% fetal bovine serum, 100 μg/mL streptomycin and 100 units/mL penicillin. PC-3 cells were cultured in F-12K Medium (ATCC, Cat. No. 30-2004) containing 10% fetal bovine serum, 100 μg/mL streptomycin and 100 units/mL penicillin.

#### Copper(I)-catalyzed alkyne-azide cycloaddition (CuAAC) labeling of glycoproteins with azido sugars

To confirm the incorporation of azido sugars into glycoproteins from Jurkat, MCF-7 and PC-3 cells, glycoproteins from cells treated with DMSO or azido sugars were labeled with TAMRA-alkyne through CuAAC labeling as in previous literature.[1]

Jurkat cells were plated at 2 x $10^5$ cells/mL and treated as mentioned before. For each sample, 10 x $10^6$ to 20 x $10^6$ cells of Jurkat cells were harvested by centrifugation, washed with DPBS and resuspended in 1% NP-40 lysis buffer (1% (w/v) IGEPAL CA-630, 150 mM NaCl, 50 mM triethanolamine (TEA), pH 7.4) containing protease inhibitor cocktail (Roche, Cat. No. 11836170001). About 2 x $10^6$ cells of PC-3 or MCF-7 cells were treated with DMSO or azido sugars as mentioned before and directly scraped into 1% NP-40 lysis buffer containing protease inhibitor cocktail. Cells were lysed in 1% NP-40 lysis buffer for 30 min at 4 °C with end-over-end rotation. Cell lysates were then centrifuged at 20,817 *g* at 4 °C for 10 min. Supernatant was used directly for CuAAC labeling or frozen in liquid nitrogen and stored at -80 °C. Protein concentrations were determined with a Pierce BCA Protein Assay Kit (Thermo Fisher Scientific, Cat. No. 23225) and BSA was used as standard.

For CuAAC labeling, 200 μg of cell lysates were diluted to a volume of 200 μL with 1% NP-40 lysis buffer, of which 188 μL was mixed with click chemistry reagent to make a final concentration of 100 μM TAMRA alkyne, 1 mM TCEP, 100 μM TBTA, 1 mM copper(II) sulfate. Lysates were then incubated at room temperature in dark for 1 hour. After the reaction, the proteins were precipitated with cold methanol and incubated at -20 °C for 2 hours. Proteins were collected by centrifugation at 20,817 *g* at 4 °C for 10 min, air-dried for 5 min at room temperature, and re-dissolved in 50 μL of 4% SDS buffer (4% (w/v) SDS, 150 mM NaCl, 50 mM TEA, pH 7.4) by sonicating for 30 min under room temperature. Lysates were then mixed with 50 μL of 2 x SDS loading dye (100 mM Tris-HCl, pH=6.8, 4% (w/v) SDS, 0.04% (w/v) Bromophenol blue, 20% (v/v) glycerol, and 20 mM DTT) and denatured at 90 °C for 5 min. Proteins were separated on a 12% TGX stain-free gel (Bio-Rad, Cat. No. 1610185). Fluorescent imaging was taken on a Typhoon FLA 9500 (GE Healthcare Life Sciences) using the protocol for TAMRA (laser wavelength: 523nm; filter: LPG), with a PMT of 400 and scanned at 25 micros per pixel. Brightness and contrast were then set to auto on ImageJ software. Gels were then activated for 5 min for the stain-free image and imaged with a ChemiDoc MP Imaging system (Bio-Rad). After imaging, gels were washed with water, fixed overnight (50% methanol and 7% acetic acid in water), washed again with water and stained with GelCode Blue stain reagent (Thermo fisher, Cat. No. 24590) according to manufacturer's instructions and then imaged with a ChemiDoc MP Imaging system (Bio-Rad).

**Cell culture for glycoproteomics and glycomics**

For analysis of azido sugar incorporation by glycoproteomics and glycomics, PC-3, MCF-7 and Jurkat cells were all plated at 2 x $10^5$ cells/mL. Cells were treated for 48 hours with 100 μM of $Ac_4$ManNAz, $Ac_4$GlcNAz or $Ac_4$GalNAz, or equal volume of DMSO as negative control and about 50 x $10^6$ cells were harvested for each sample.[2] For Jurkat cells, cells were separated from cell culture medium by centrifugation and washed with DPBS (Sigma-Aldrich, Cat. No. D8537-500ML). For PC-3 and MCF-7 cells, cells were lifted from plate by incubation with DPBS containing 1 mM of ethylenediaminetetraacetic acid (EDTA) at 37 °C and quenched with cell culture medium. Cells were separated by centrifugation, washed with DPBS, frozen in liquid nitrogen and stored at -80°C.

**Evaluation of reduction of azide functional group on metabolically incorporated glycans by DTT**

To determine whether incubation with DTT will lead to decrease in available azide, 200 μg of cell lysates were diluted to a total volume of 90 μL with 1% NP-40 lysis buffer containing protease inhibitor cocktail. Either 10 μL of 1 M DTT or 1% NP-40 lysis buffer was added to the lysates and lysates were then incubated overnight at 4 °C with end-over-end rotation. Because DTT will interfere with CuAAC labeling, proteins were isolated from the solution using methanol-chloroform precipitation. Briefly, 300 μL of water, 400 μL of methanol and 100 μL of chloroform were added to 100 μL of cell lysates, vortex, and centrifuged at 13,000 *g* at 4 °C for 10 min. Precipitant was then washed with 400 μL of methanol and centrifuged at 13,000 *g* at 4 °C for 10 min.[3] Pellet was air-dried for 5 min at room temperature and re-dissolved in 200 μL of 1% NP-40 lysis buffer containing 0.5% (w/v) SDS by sonicating for 30 min under room temperature. Undissolved proteins were removed by centrifugation and the supernatant was labeled with TAMRA-alkyne as above.

**HPLC analysis of DMB-derivatized sialic acids released from cells**

To determine the generation of SiaNAz, 2 million of MCF7 or PC-3 cells, or 3 to 8 million of Jurkat cells were treated with DMSO or azido sugars as mentioned before. Jurkat cells were harvested by centrifugation and washed with DPBS. MCF7 and PC-3 cells were released from plates by 1 mM EDTA in DPBS, quenched with medium and washed with DPBS. Cells were frozen in liquid nitrogen and stored at -80 ºC.

Methods for cell lysis and DMB-labeling were adapted from a previous study.[4] Cells were resuspended in 1 ml of hypotonic lysis buffer (10 mM Tris, pH = 7.3, 10 mM $MgCl_2$, 1 mM EDTA, 1mM EGTA) with protease inhibitor cocktail (Sigma, Cat. No. 11836170001), and incubated on ice for 15 min. Cells were extruded through a 25 gauge needle for 5 min and centrifuged at 1,000 *g* at 4 ºC for 30 or 45 min to remove the insoluble debris. Concentration of supernatant was determined with a Pierce BCA Protein Assay Kit (Thermo Fisher Scientific, Cat. No. 23225) and BSA was used as standard. For each cell line, the same amount of lysates (200 μg or 400 μg) were frozen in liquid nitrogen and dried with a SAVANT SC210A SpeedVac Concentrator (Thermo Fisher Scientific).

The dried samples were resuspended with 100 μL of 2 M acetic acid, vortexed, and centrifuged at room temperature at 17,000 *g* for 5 sec. The supernatant was incubated at 80 ºC for 2 h to release the sialic acid. After acid hydrolysis, 80 μL of DMB reaction solution (0.75 M β-mercaptoethanol, 18 mM sodium thiosulfate, 1.4 M acetic acid, 7 mM DMB) was added, and incubated at 50 ºC for 2 hours. Reaction was quenched by addition of 20 μL of 0.2 M NaOH, centrifuged at 17,000 *g* at room temperature for 5 min and filtered through a 0.2 μm syringe filter (Millipore, SLFGR04NL). DMB-derivatized sialic acid was stored at -20 ºC in dark for up to a week before further analysis.

For the HPLC analysis, 20 μL of DMB-labeled samples was diluted in 180 μL of water and 20 μL of diluted samples were injected. DMB-derivatized sialic acids were separated by HPLC with an isocratic program (86% water, 8% acetonitrile, 6% methanol for 40 min). DMB-derivatizing reactions with Neu5Ac or no sialic acid were used as standard for DMB-Neu5Ac or negative control, respectively. SiaNAz released from Jurkat cells treated with $Ac_4$ManNAz and DMB-derivatized was purified by HPLC, dried by SpeedVac Concentrator and confirmed by LC-MS ($C_{18}H_{23}N_6O_9$ $[M+H]^+$, theoretical m/z: 467.15210, measured m/z: 467.15242). DMB-SiaNAz was stored at -20 ºC in dark and used as a standard for HPLC.

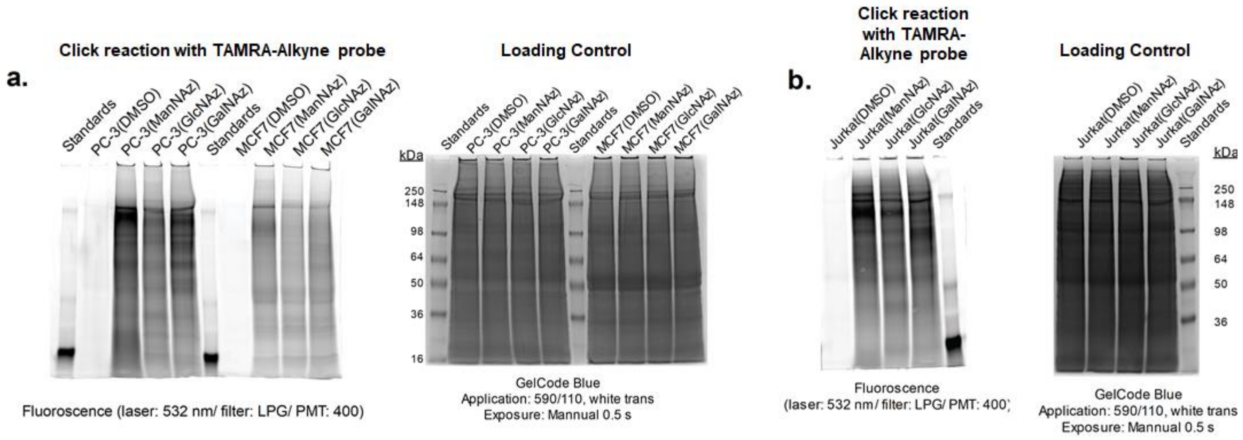


**Supp Fig S1:** In-gel imaging showing the incorporation of azido sugars on the cellular glycoprotein upon treatment with non-natural monosaccharides bearing azido functional group; **a.** PC-3 and MCF-7 cells treated with DMSO, Ac4ManNAz, Ac4GlcNAz, and Ac4GalNAz; **b.** Jurkat cells treated with DMSO, Ac4ManNAz, Ac4GlcNAz, and Ac4GalNAz.

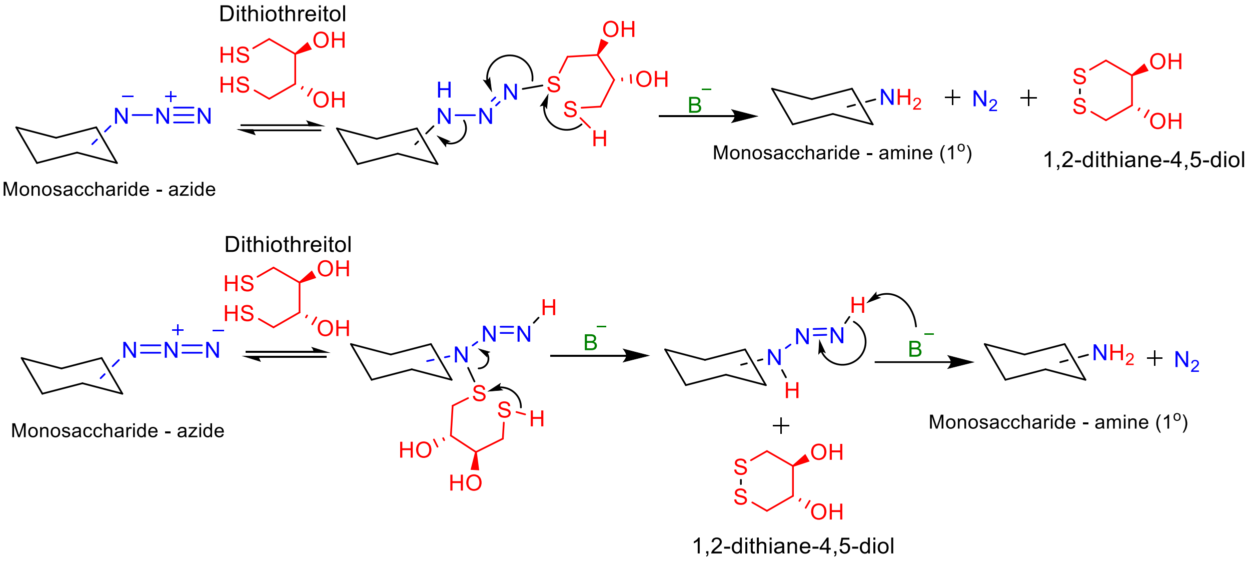


**Supp Fig S2:** Proposed mechanism for the reduction of azide functional group on the non-natural glycans by the treatment with DTT.[5]

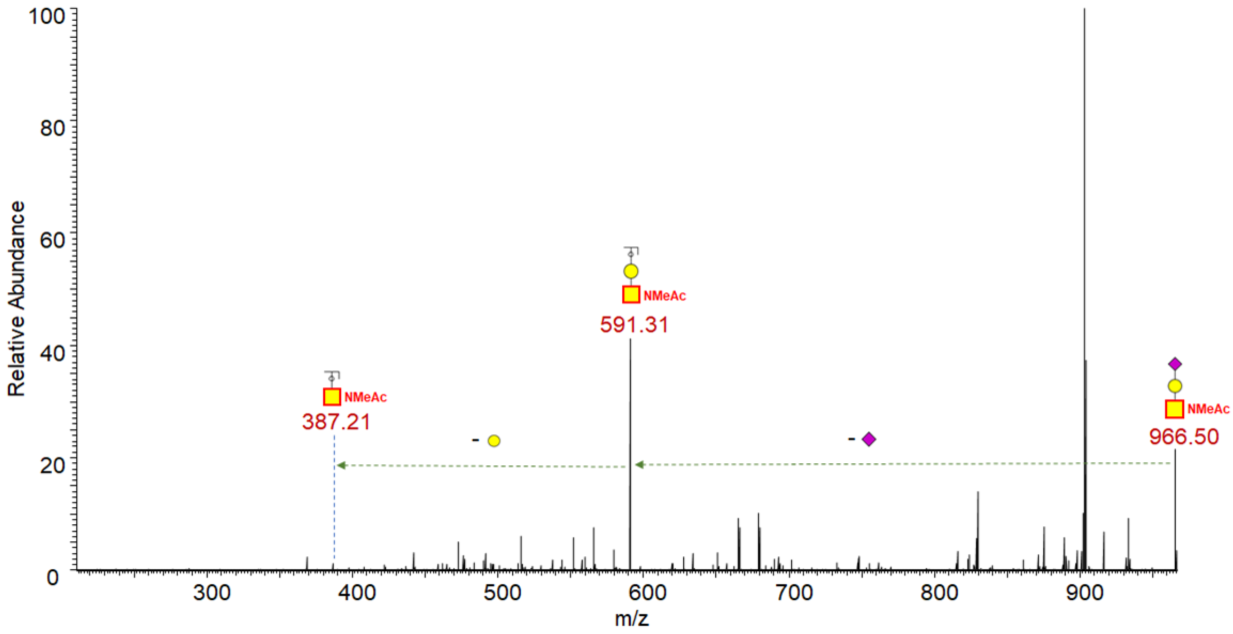


**Supp Fig S3:** Glycomics analysis of metabolically incorporated azide bearing O-glycans from $Ac_4GalNAz$ treated PC-3 cells and ESI-$MS^2$ fragmentation of the acetylated-permethylated labelled O-glycan 966.50 (M+Na) showing the characteristic fragments and neutral losses.

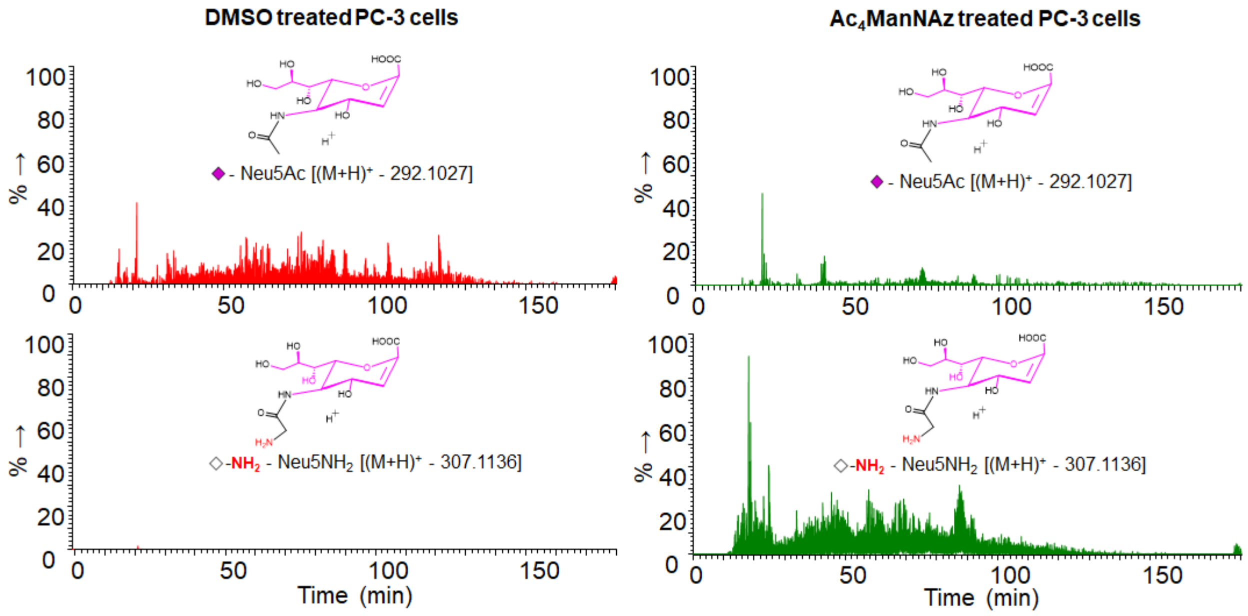


**Supp Fig S4:** Extracted ion chromatogram of DMSO (vehicle) and $Ac_4ManNAz$ treated enriched PC-3 cell digest showing sialic acid and non-natural azidoacetyl sialic acid (Neu5Az) in its reduced amine form ($Neu5NH_2$).

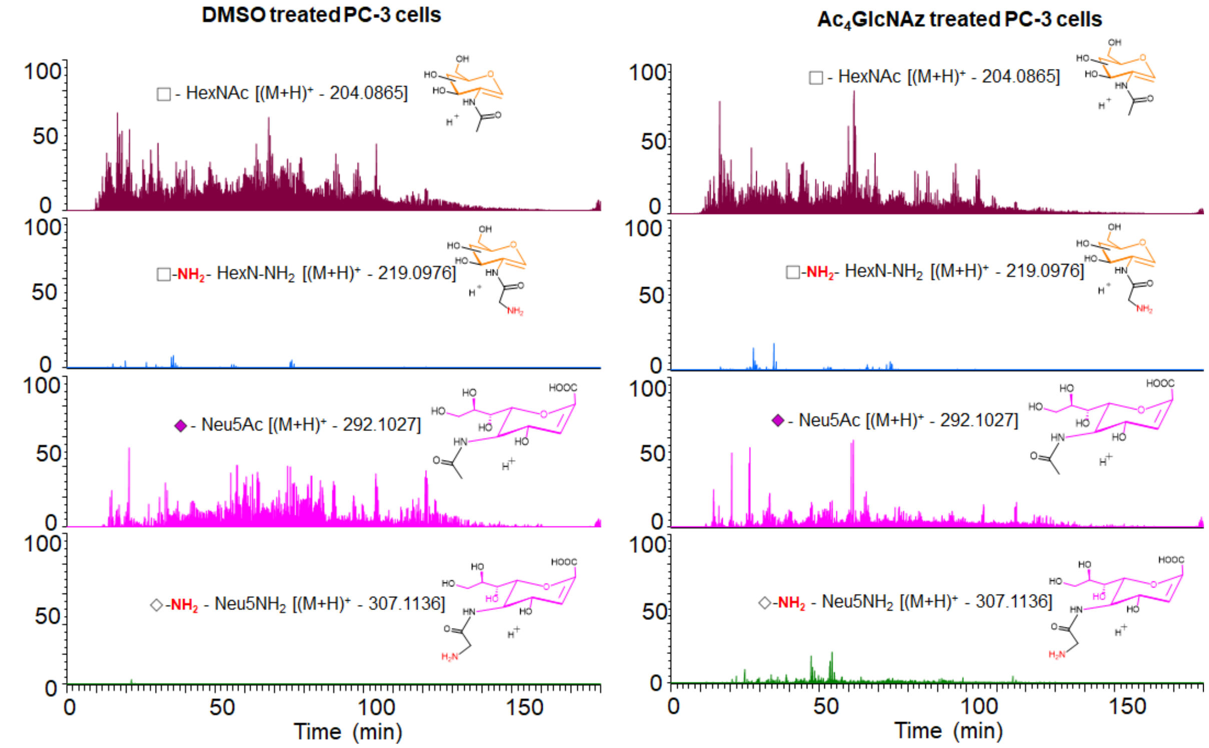


**Supp Fig S5:** Extracted ion chromatogram of DMSO (vehicle) and $Ac_4GlcNAz$ treated enriched PC-3 cell digest showing HexNAc (can be both GalNAc and GlcNAc), non-natural azidoacetyl HexNAc (HexNAz) in its reduced amine form ($HexN-NH_2$), sialic acid and non-natural azidoacetyl sialic acid (Neu5Az) in its reduced amine form ($Neu5NH_2$).

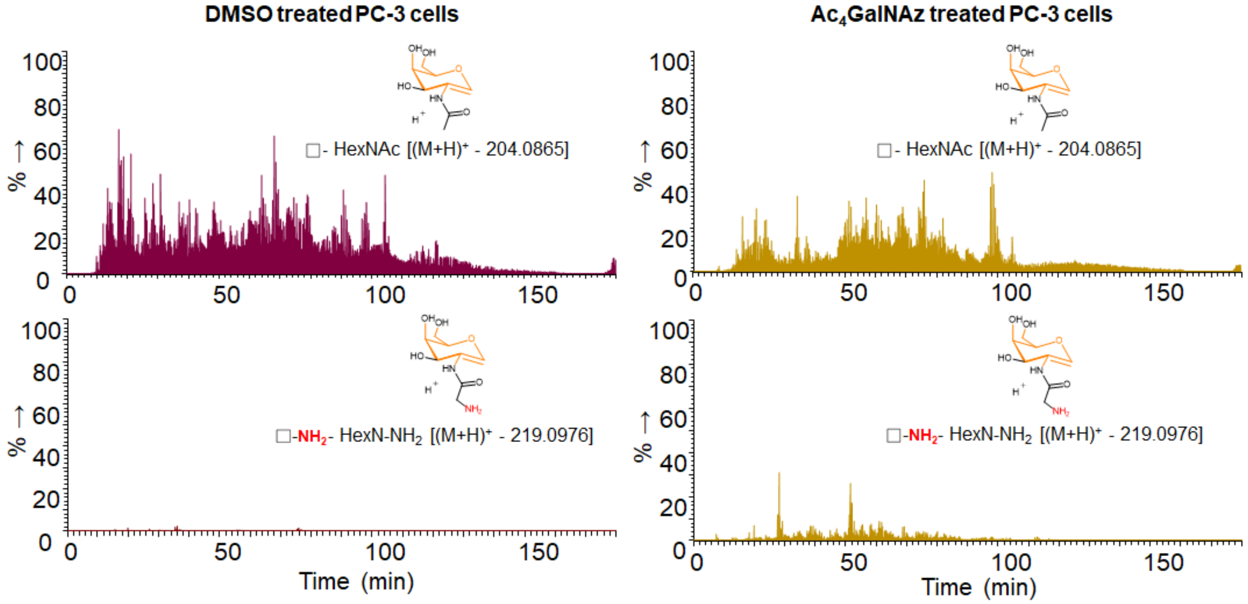


**Supp Fig S6:** Extracted ion chromatogram of DMSO (vehicle) and $Ac_4GalNAz$ treated enriched PC-3 cell digest showing HexNAc (can be both GalNAc and GlcNAc) and non-natural azidoacetyl HexNAc (HexNAz) in its reduced amine form ($HexN-NH_2$).

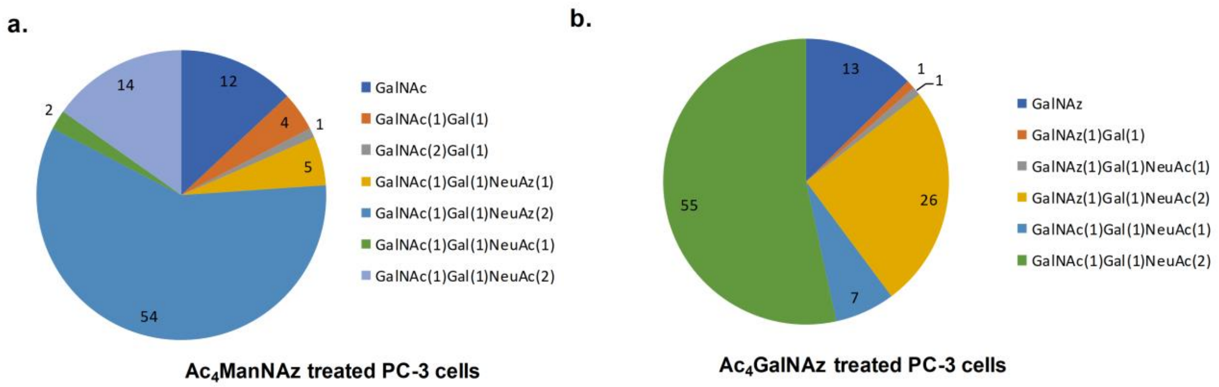


**Supp Fig S7:** Natural and non-natural glycans containing O-glycopeptides detected on PC-3 cells by the treatment with a. $Ac_4$ManNAz treated cells; b. $Ac_4$GalNAz treated cells.

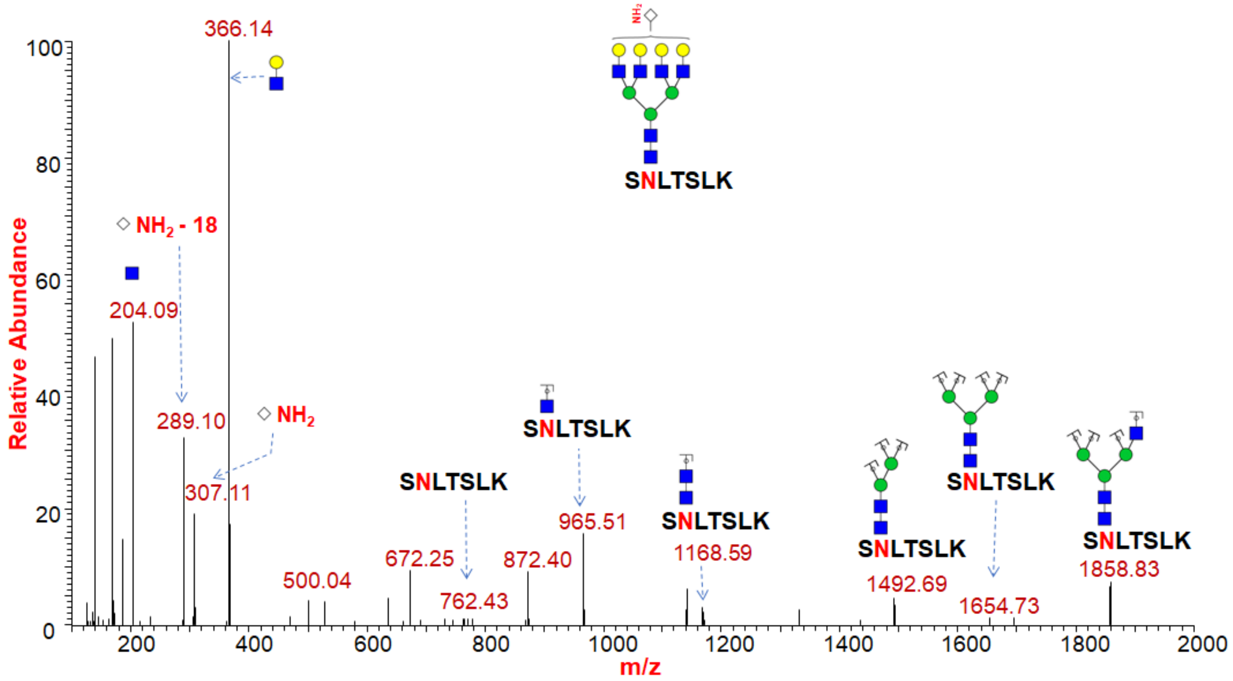


**Supp Fig S8:** HCD $MS^2$ spectrum of glycopeptide SNLTSLK from Centrosomal protein of 68 kDa (Q76N32) protein from $Ac_4$ManNAz treated PC-3 cells showing Neu5$NH_2$ incorporation on N-glycans (Nn49) by the presence of oxonium ion (m/z 307.11 & 289.10) and neutral losses (m/z 306.10).

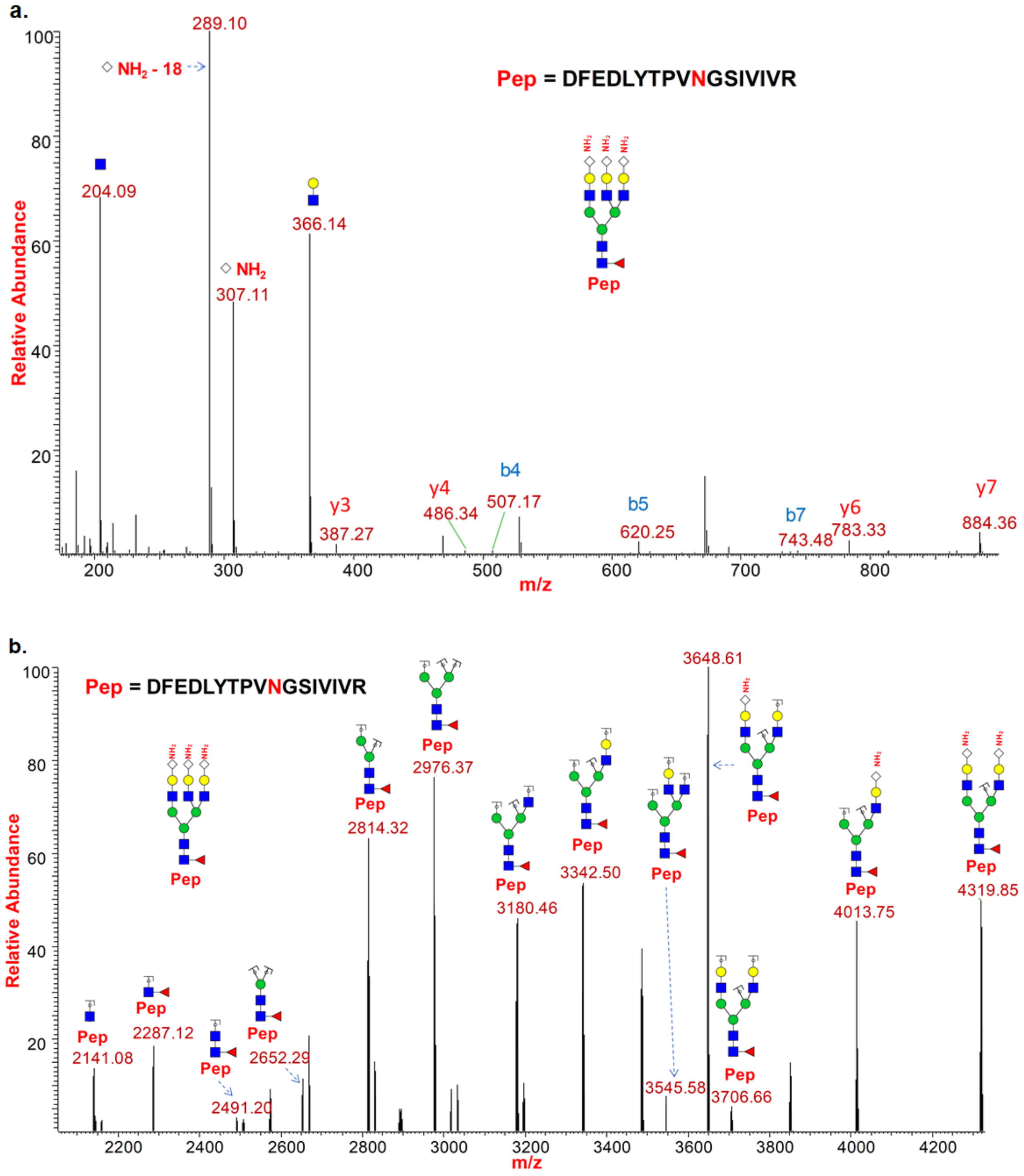


**Supp Fig S9:** HCD (a.) and CID (b.) $MS^2$ spectrum of glycopeptide DFEDLYTPVNGSIVIVR from Transferrin receptor protein (P02786) protein from $Ac_4ManNAz$ treated PC-3 cells showing $Neu5NH_2$ incorporation on N-glycans (Nn48) by the presence of oxonium ion (m/z 307.11 & 289.10) and neutral losses (m/z 306.10).

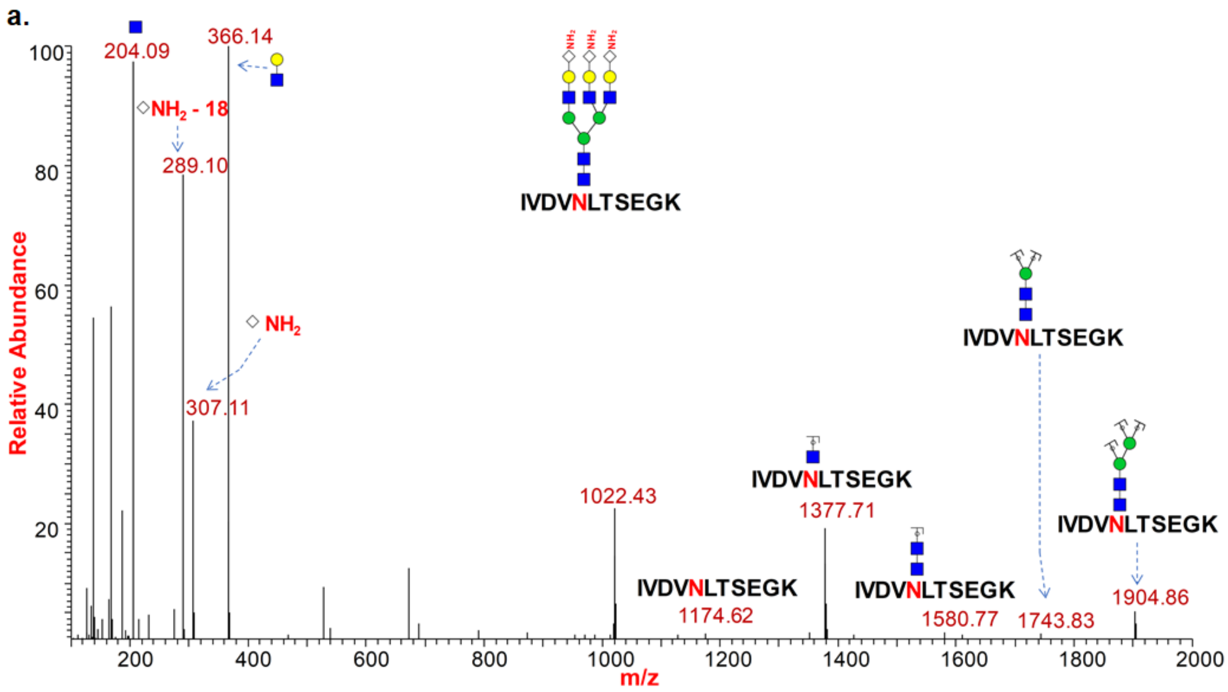


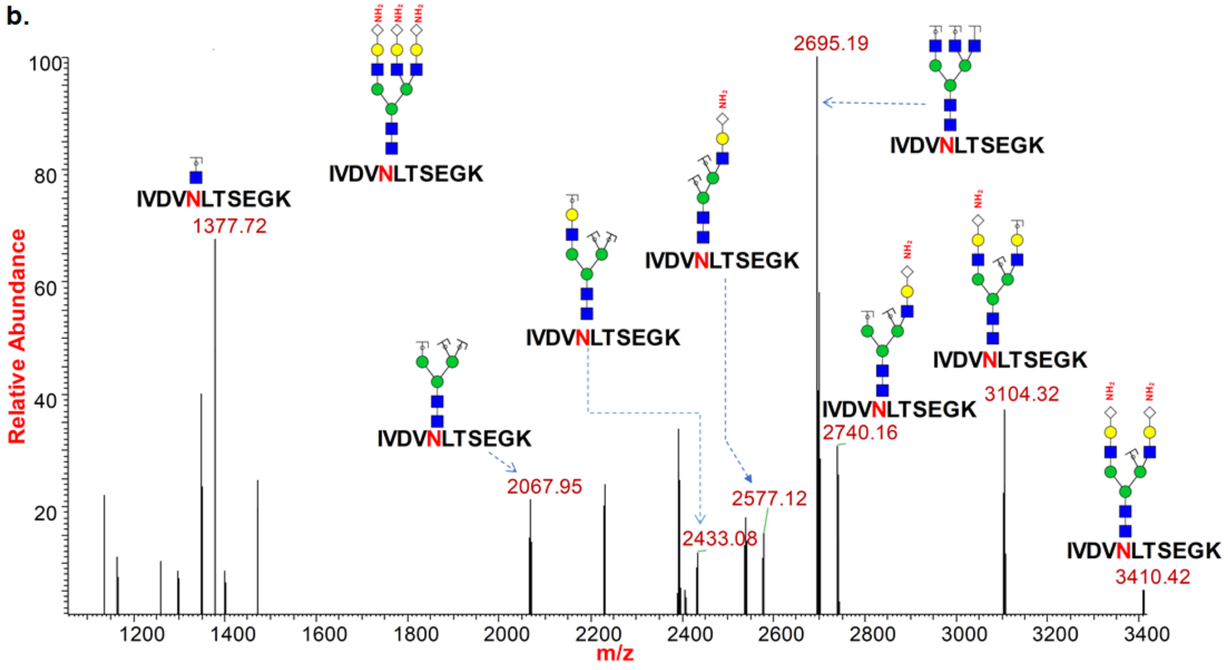


**Supp Fig S10:** HCD (a.) and CID (b.) $MS^2$ spectrum of glycopeptide IVDVNLTSEGK from Transmembrane 9 superfamily member 3 (Q9HD45) protein from $Ac_4ManNAz$ treated PC-3 cells showing $Neu5NH_2$ incorporation on N-glycans (Nn47) by the presence of oxonium ion (m/z 307.11 & 289.10) and neutral losses (m/z 306.10).

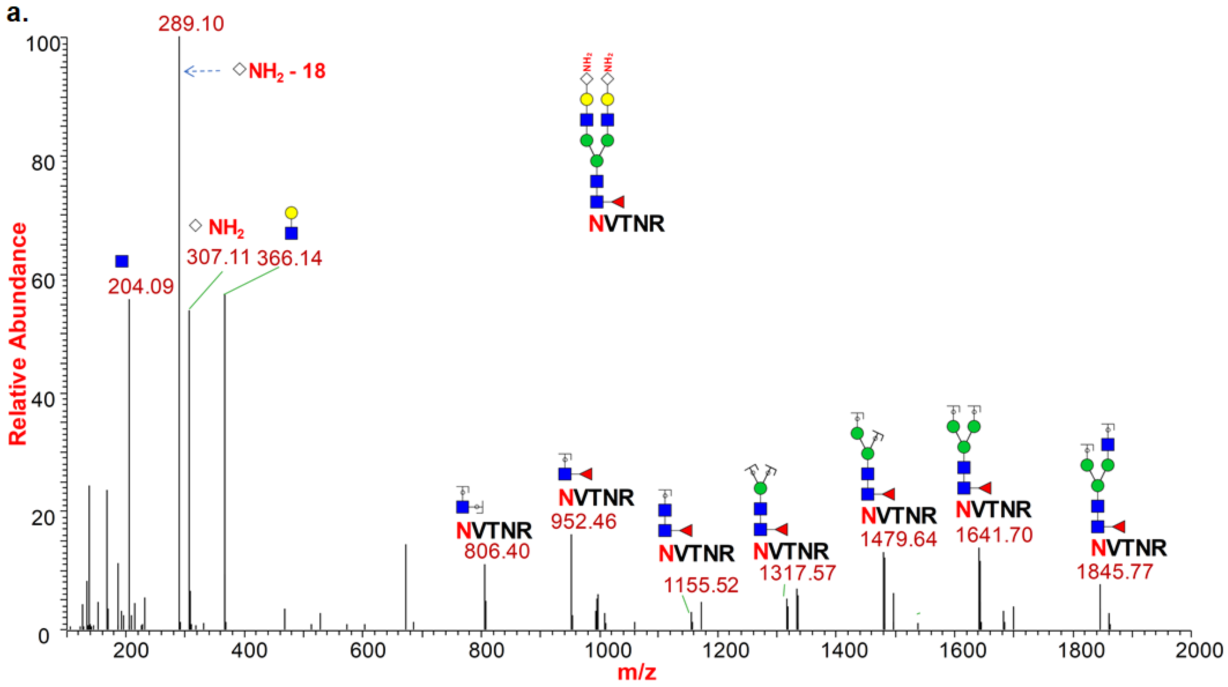


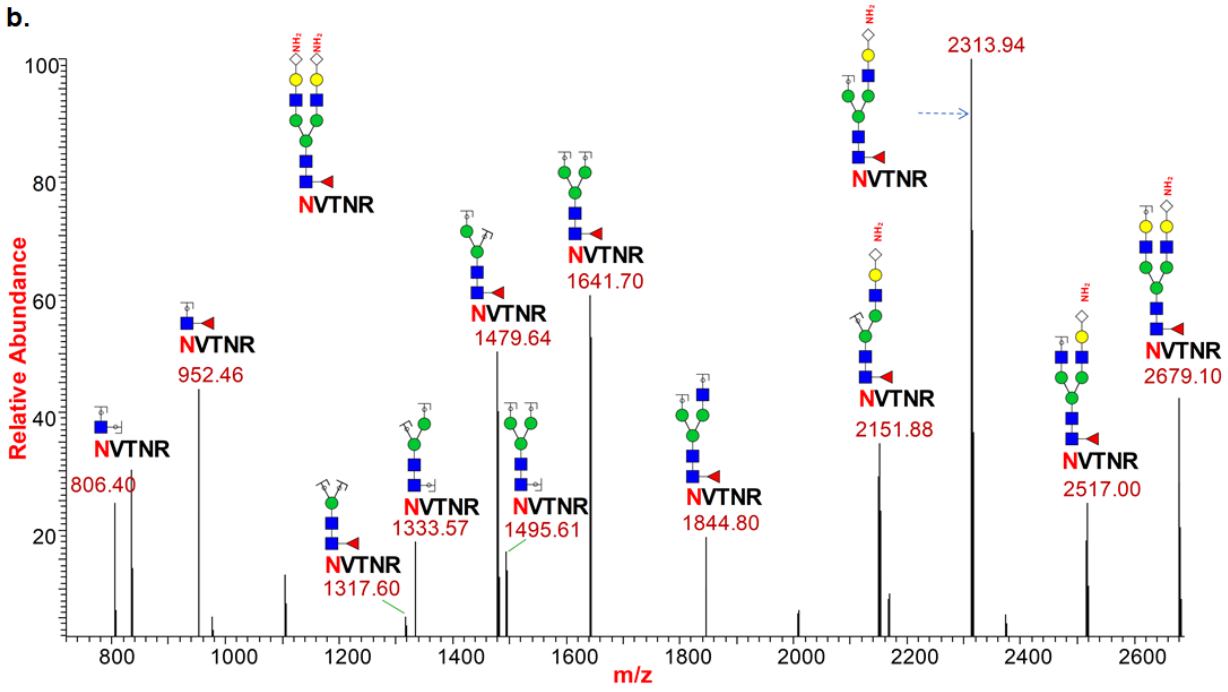


**Supp Fig S11:** HCD (a.) and CID (b.) $MS^2$ spectrum of glycopeptide NVTNR from HUMAN Integrin beta-1 (P05556) protein from $Ac_4ManNAz$ treated PC-3 cells showing $Neu5NH_2$ incorporation on N-glycans (Nn46) by the presence of oxonium ion (m/z 307.11 & 289.10) and neutral losses (m/z 306.10).

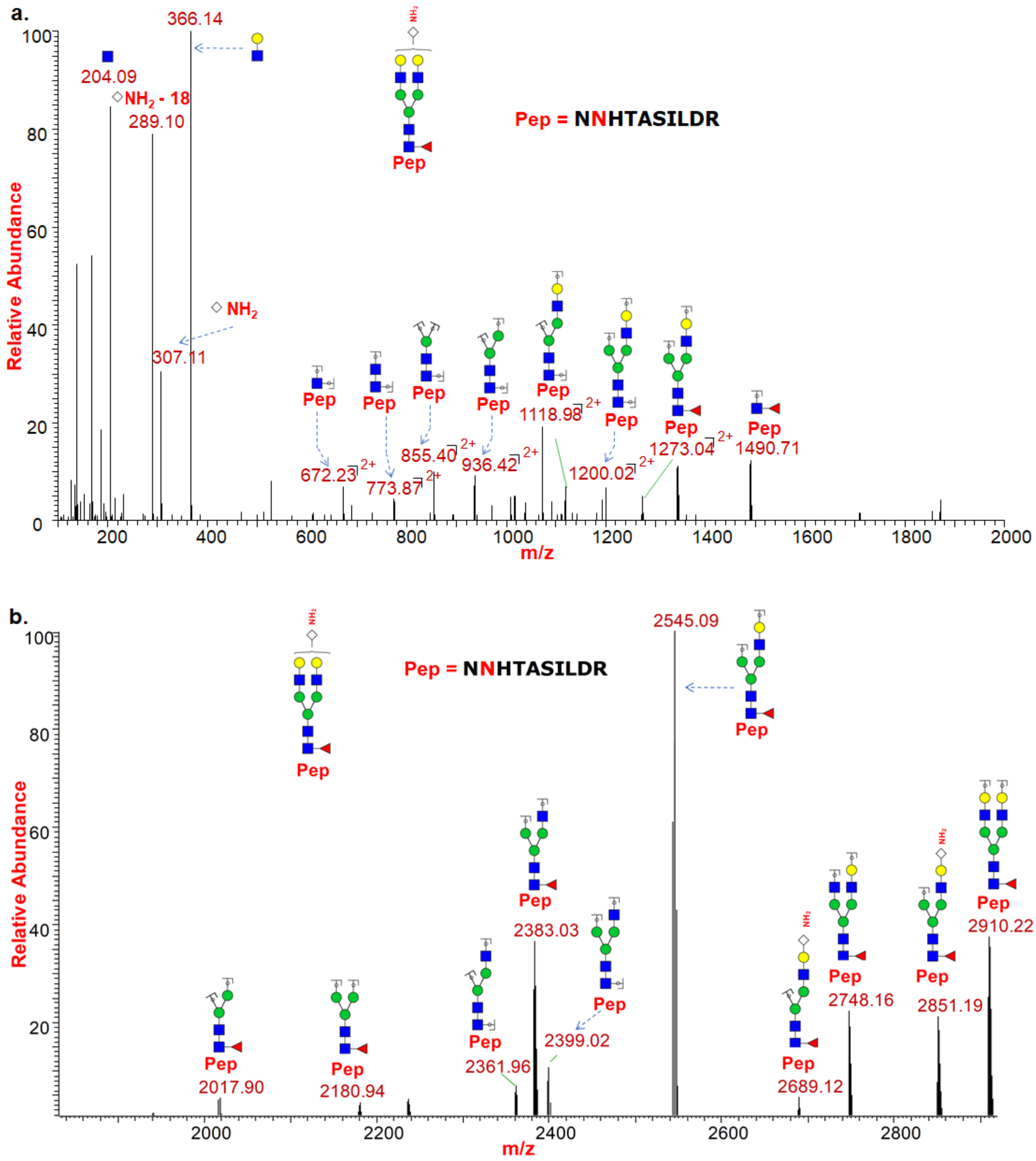


**Supp Fig S12:** HCD (a.) and CID (b.) $MS^2$ spectrum of glycopeptide NNHTASILDR from CD63 antigen (P08962) protein from $Ac_4ManNAz$ treated PC-3 cells showing $Neu5NH_2$ incorporation on N-glycans (Nn44) by the presence of oxonium ion (m/z 307.11 & 289.10) and neutral losses (m/z 306.10).

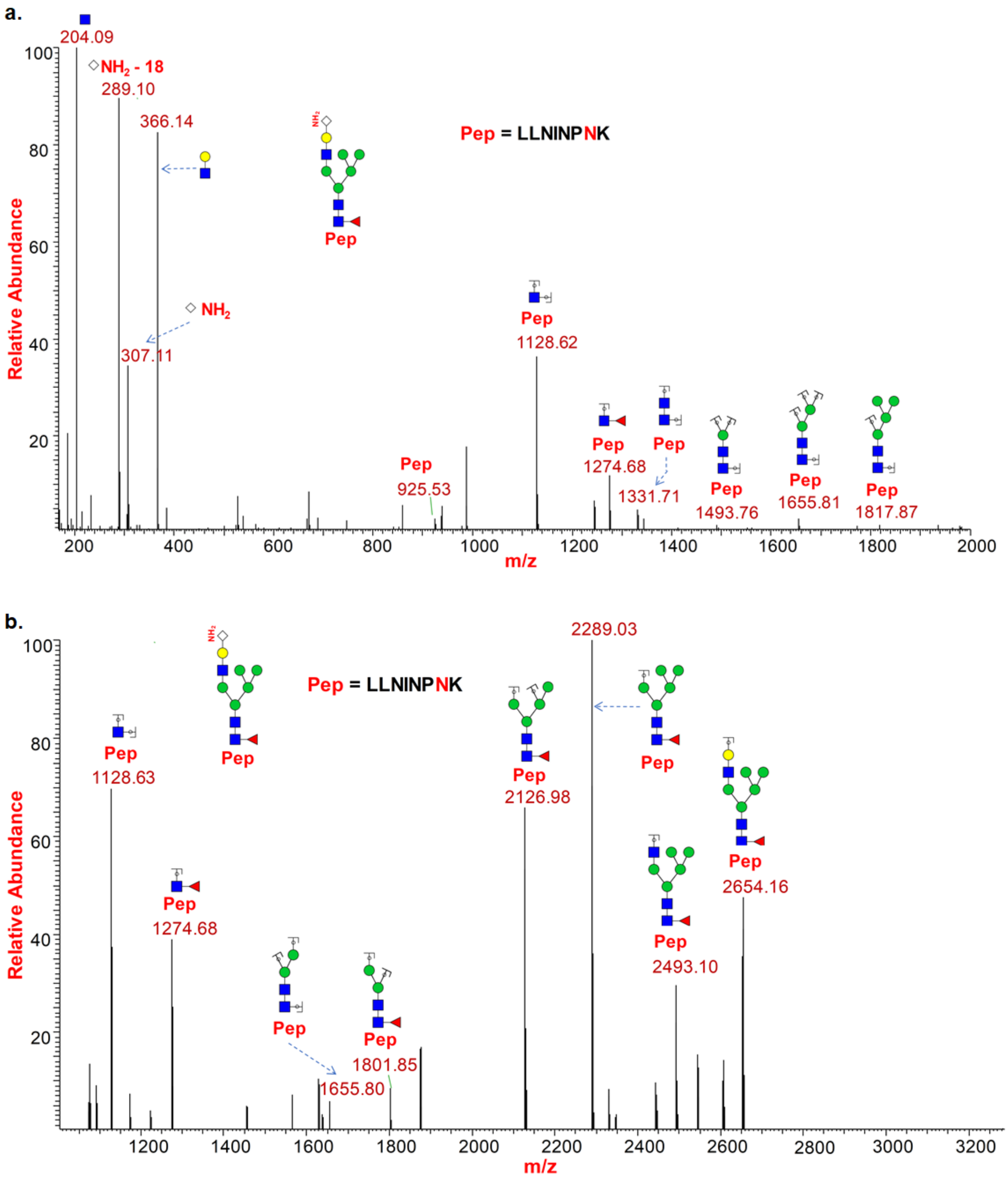


**Supp Fig S13:** HCD (a.) and CID (b.) MS[2] spectrum of glycopeptide LLNINPNK from Lysosome-associated membrane glycoprotein 1 (P11279) protein from $Ac_4ManNAz$ treated PC-3 cells showing $Neu5NH_2$ incorporation on N-glycans (Nn43) by the presence of oxonium ion (m/z 307.11 & 289.10) and neutral losses (m/z 306.10).

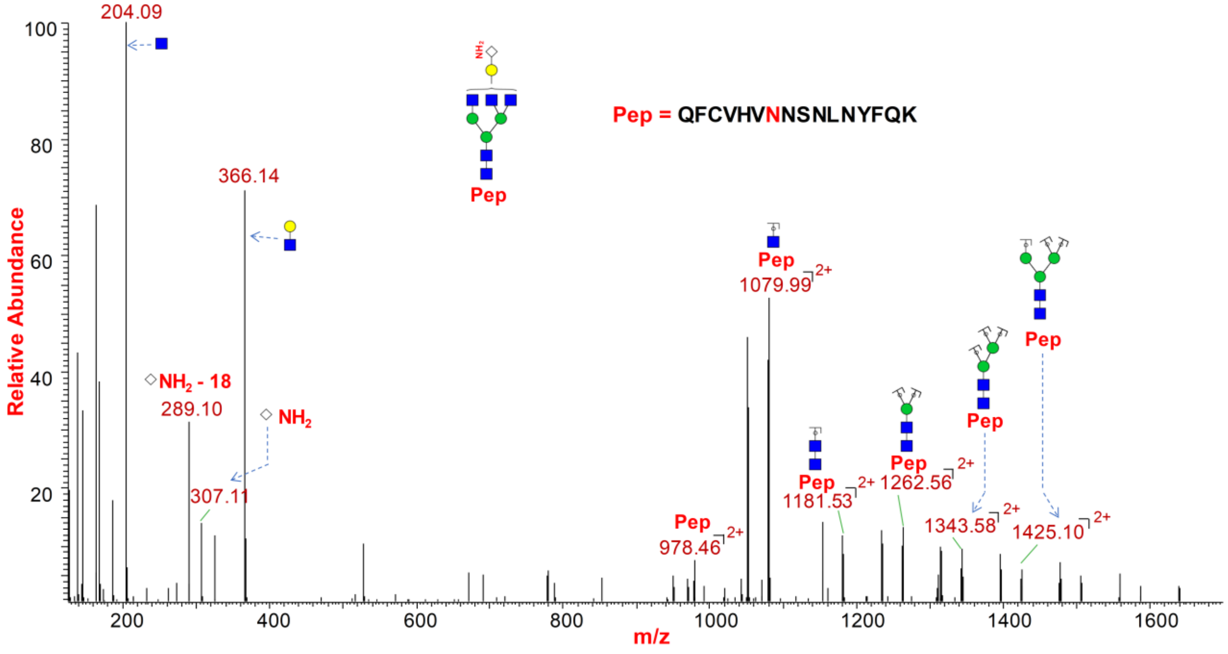


**Supp Fig S14:** HCD MS$^2$ spectrum of glycopeptide QFCVHVNNSNLNYFQK from Tectonic-3 (Q6NUS6) protein from $Ac_4ManNAz$ treated PC-3 cells showing $Neu5NH_2$ incorporation on N-glycans (Nn42) by the presence of oxonium ion (m/z 307.11 & 289.10) and neutral losses (m/z 306.10).

**a.**

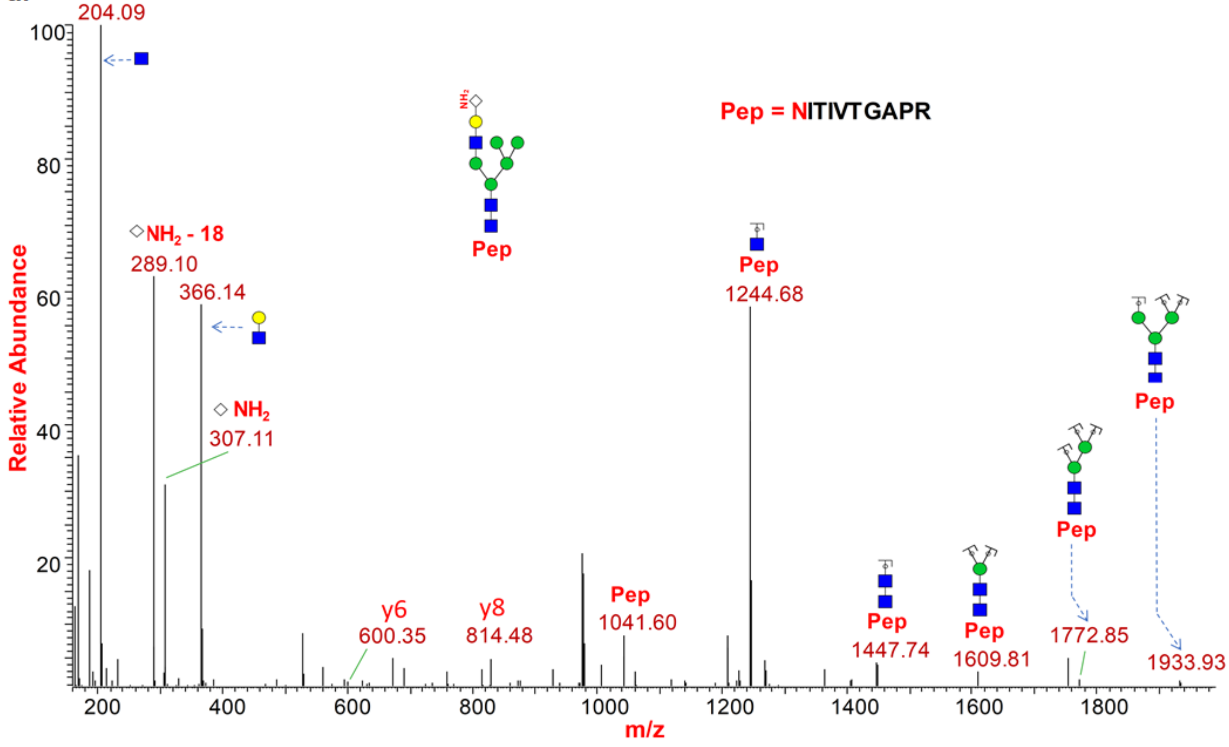

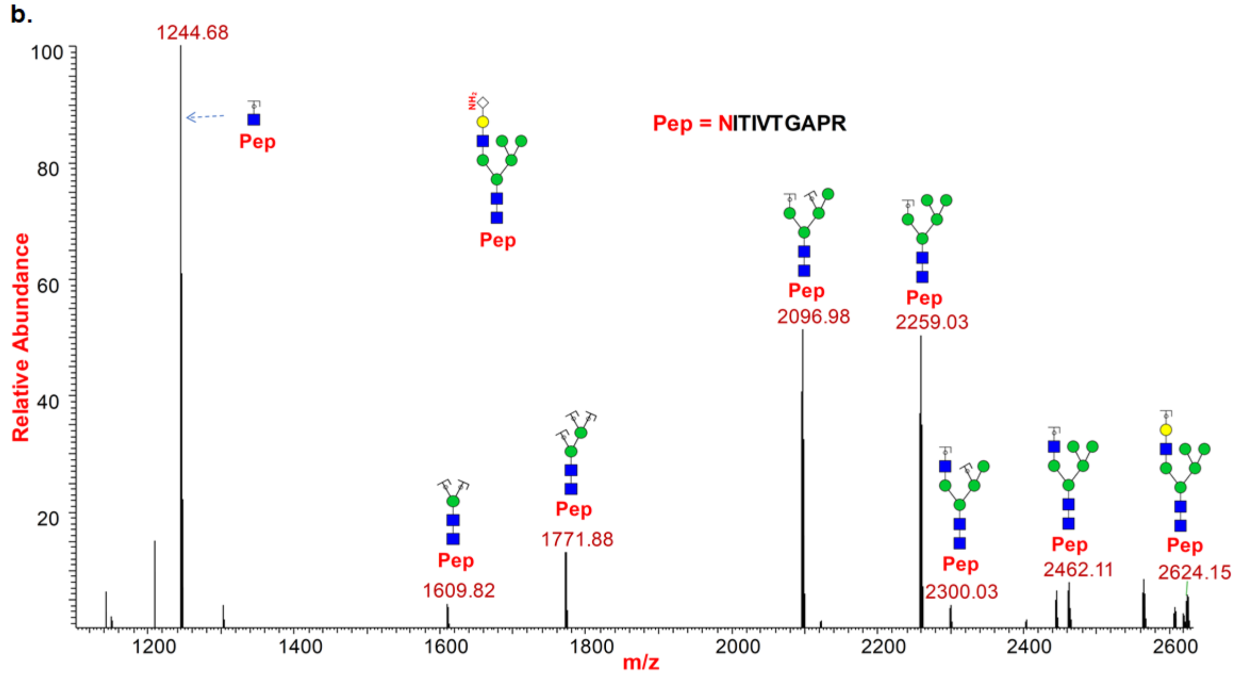


**Supp Fig S15:** HCD (a.) and CID (b.) $MS^2$ spectrum of glycopeptide NITIVTGAPR from Integrin alpha-3 (P26006) protein from $Ac_4ManNAz$ treated PC-3 cells showing $Neu5NH_2$ incorporation on N-glycans (Nn40) by the presence of oxonium ion (m/z 307.11 & 289.10) and neutral losses (m/z 306.10).

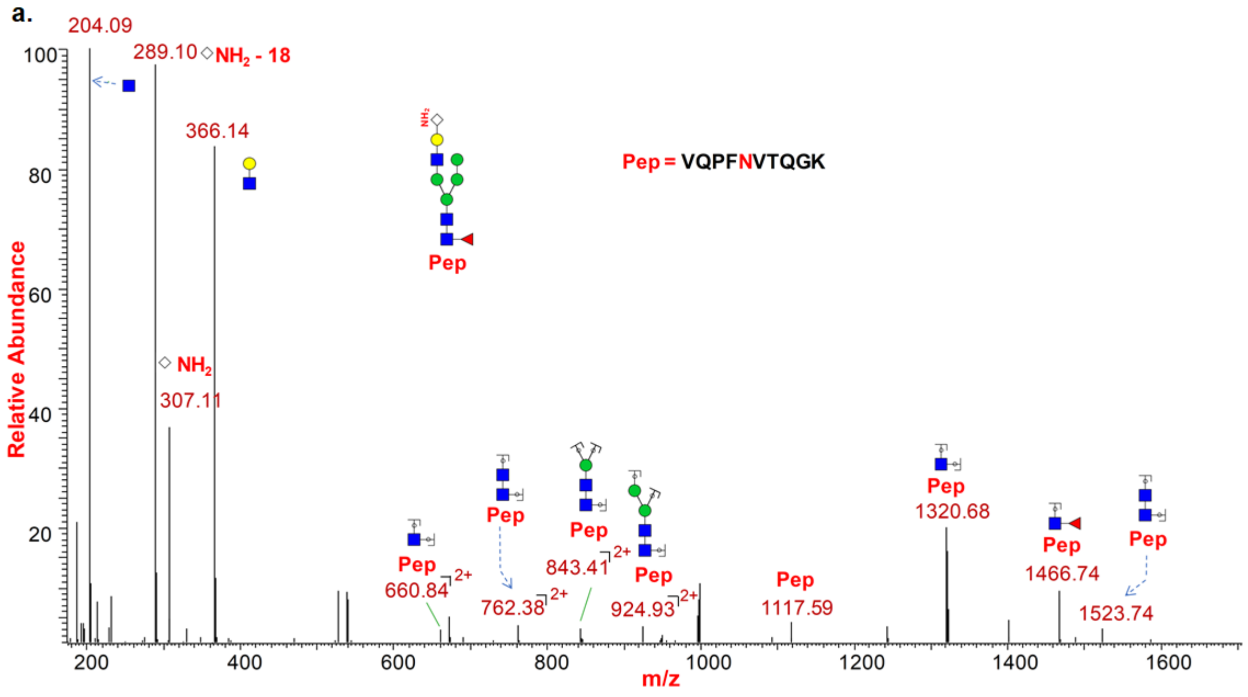

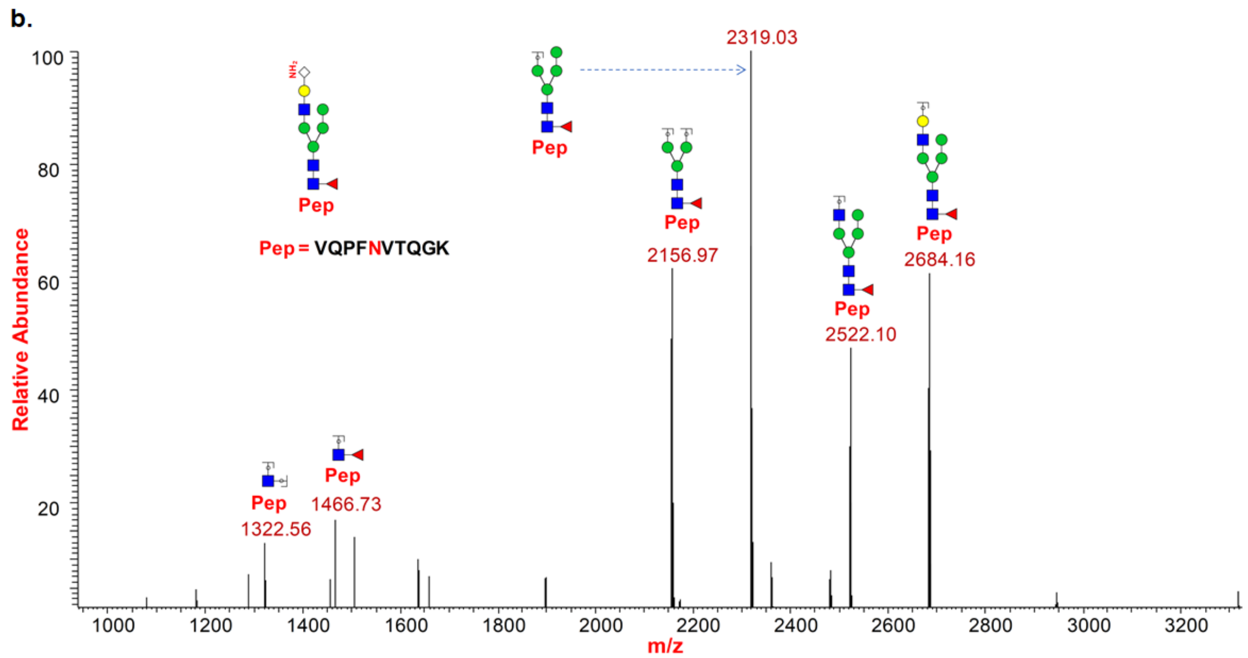


**Supp Fig S16:** HCD (a.) and CID (b.) $MS^2$ spectrum of glycopeptide VQPFNVTQGK from Lysosome-associated membrane glycoprotein 2 (P13473) protein from $Ac_4ManNAz$ treated PC-3 cells showing $Neu5NH_2$ incorporation on N-glycans (Nn39) by the presence of oxonium ion (m/z 307.11 & 289.10) and neutral losses (m/z 306.10).

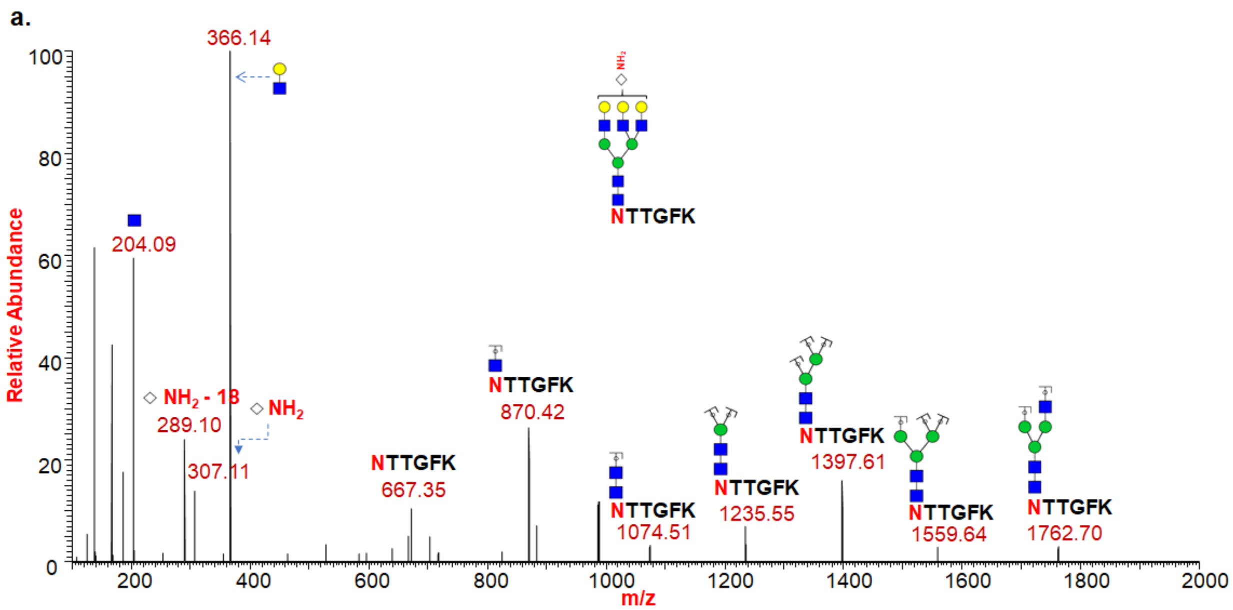

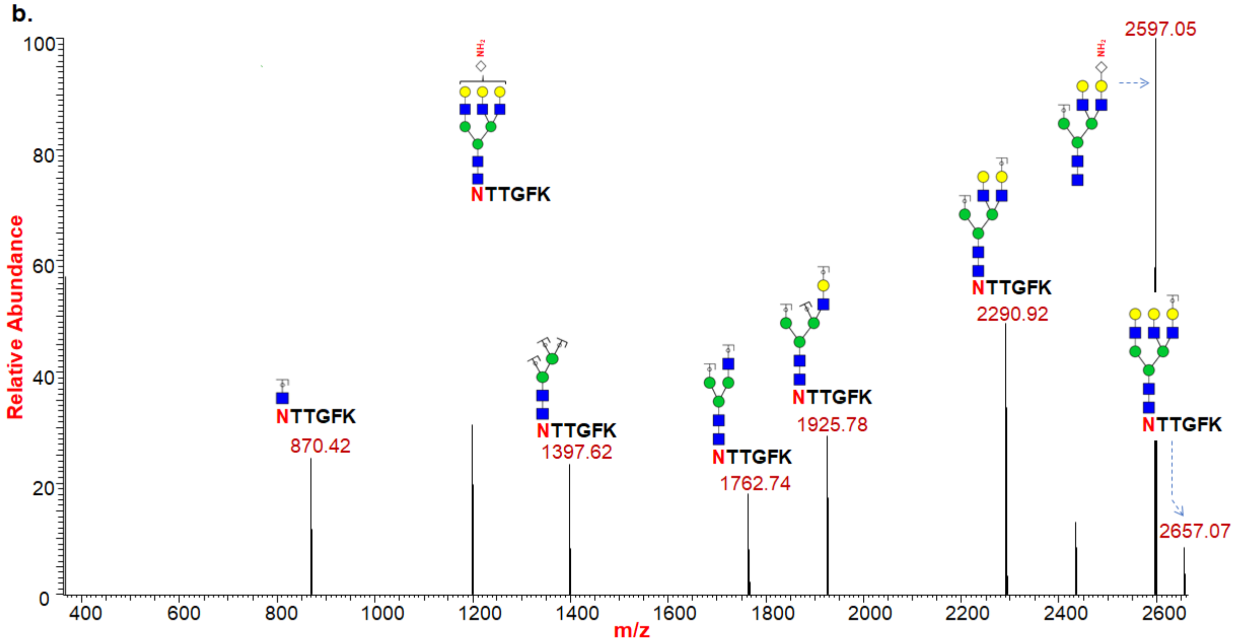


**Supp Fig S17:** HCD (a.) and CID (b.) MS[2] spectrum of glycopeptide NTTGFK from Laminin alpha-4 (Q16363) protein from Ac4ManNAz treated Jurkat cells showing Neu5NH2 incorporation on N-glycans (Nn39) by the presence of oxonium ion (m/z 307.11 & 289.10) and neutral losses (m/z 306.10).

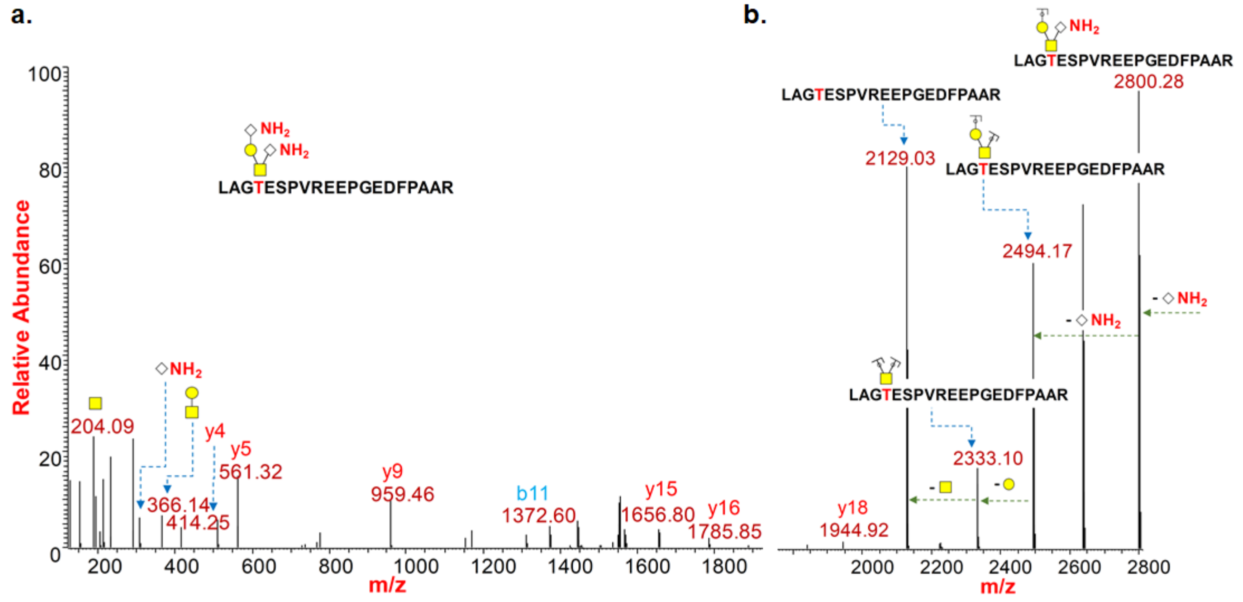


**Supp Fig S18:** HCD (a.) and CID (b.) MS[2] spectrum of glycopeptide LAGTESPVREEPGEDFPAAR from Transferrin receptor protein 1 (P02786) protein from Ac4ManNAz treated PC-3 cells showing Neu5NH2incorporation on core-1 disialylated O-glycans (On10) by the presence of oxonium ion (m/z 307.11 & 289.10) and neutral losses (m/z 306.10).

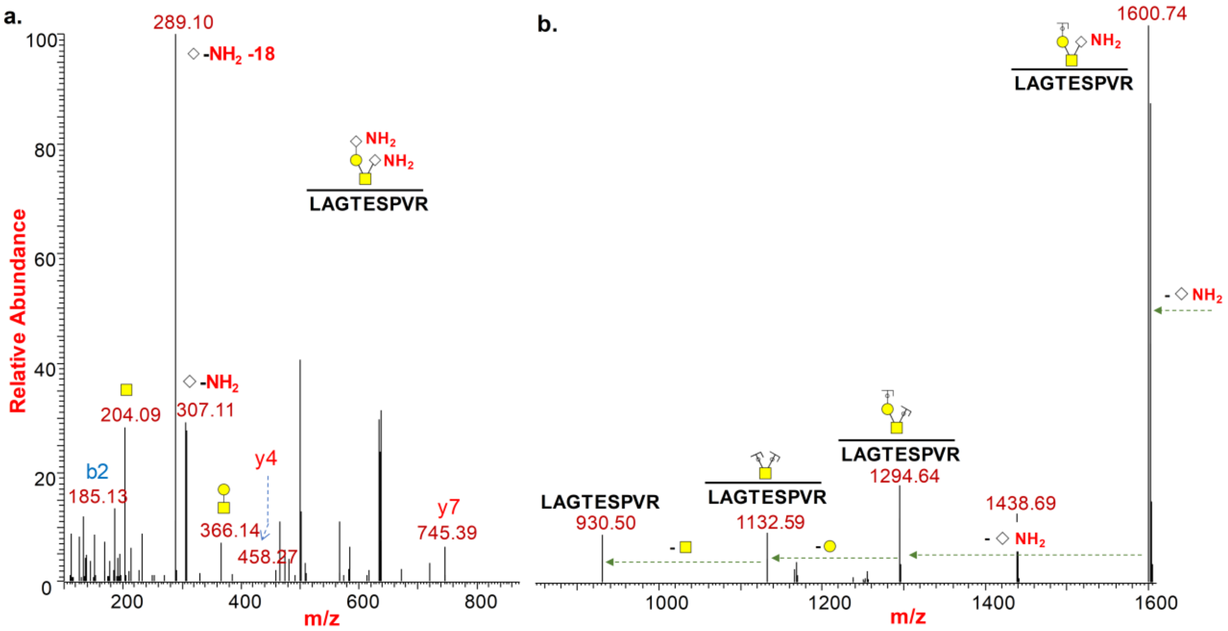


**Supp Fig S19:** HCD (a.) and CID (b.) $MS^2$ spectrum of glycopeptide LAGTESPVR from Transferrin receptor protein 1 (P02786) protein from $Ac_4ManNAz$ treated MCF-7 cells showing $Neu5NH_2$incorporation on core-1 disialylated O-glycans (On10) by the presence of oxonium ion (m/z 307.11 & 289.10) and neutral losses (m/z 306.10).

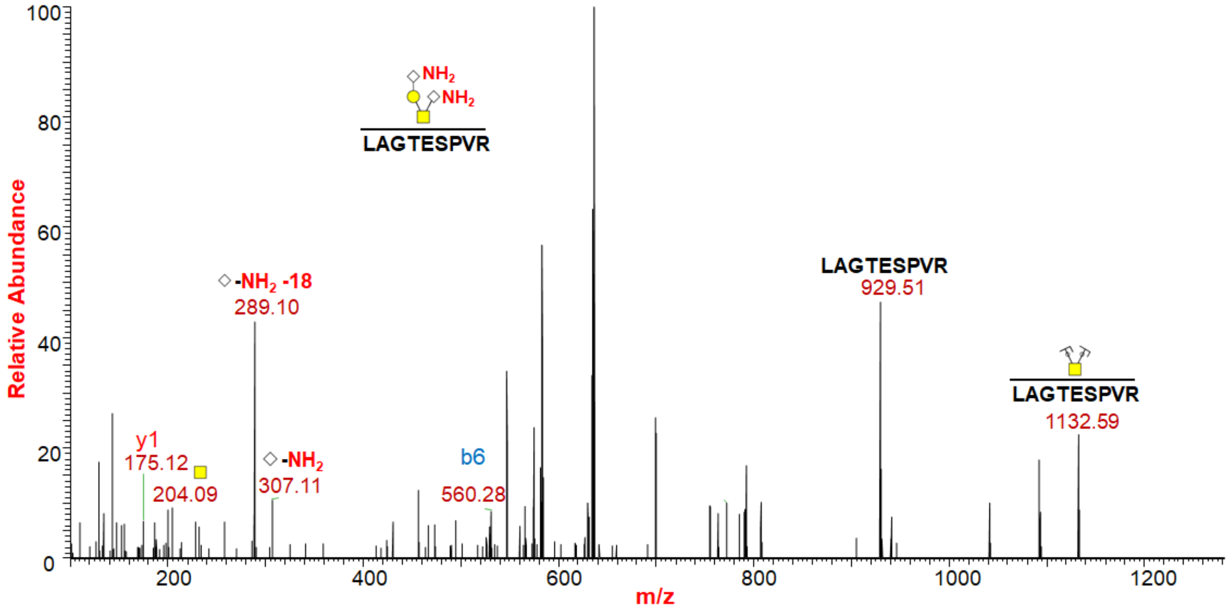


**Supp Fig S20:** HCD $MS^2$ spectrum of glycopeptide LAGTESPVR from Transferrin receptor protein 1 (P02786) protein from $Ac_4ManNAz$ treated Jurkat cells showing $Neu5NH_2$incorporation on core-1 disialylated O-glycans (On10) by the presence of oxonium ion (m/z 307.11 & 289.10).

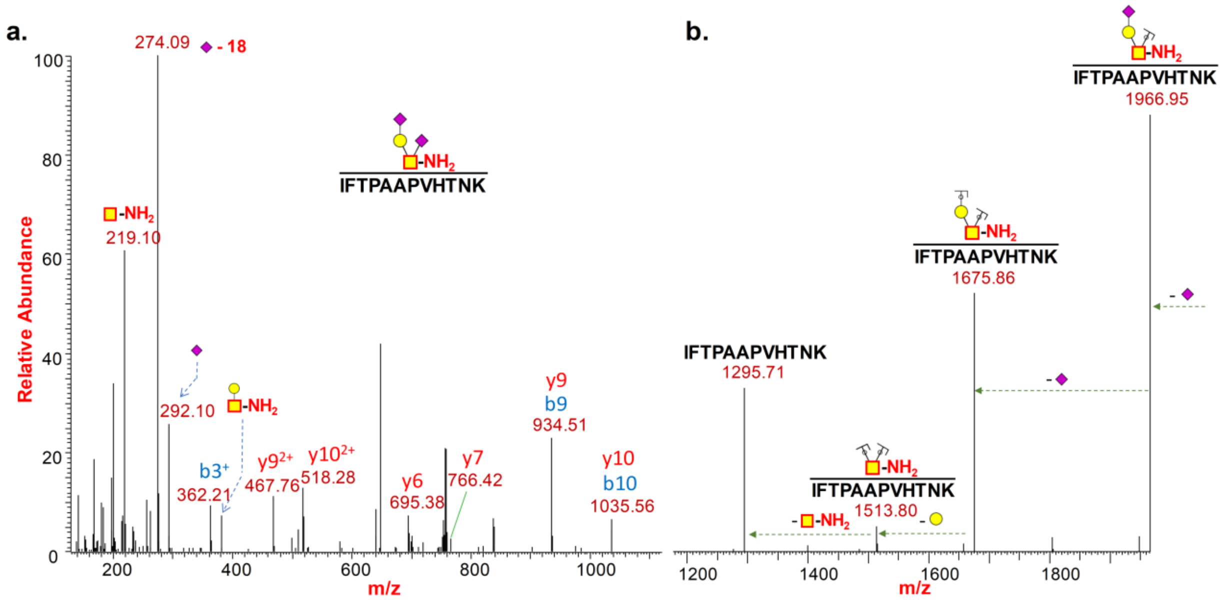


**Supp Fig S21:** HCD (a.) and CID (b.) $MS^2$ spectrum of glycopeptide IFTPAAPVHTNK from Transmembrane protein 165 (Q9HC07) protein from $Ac_4GalNAz$ treated PC-3 cells showing GalN-$NH_2$ incorporation on core-1 disialylated O-glycans (On7) by the presence of oxonium ion (m/z 219.10) and neutral losses (m/z 218.09).

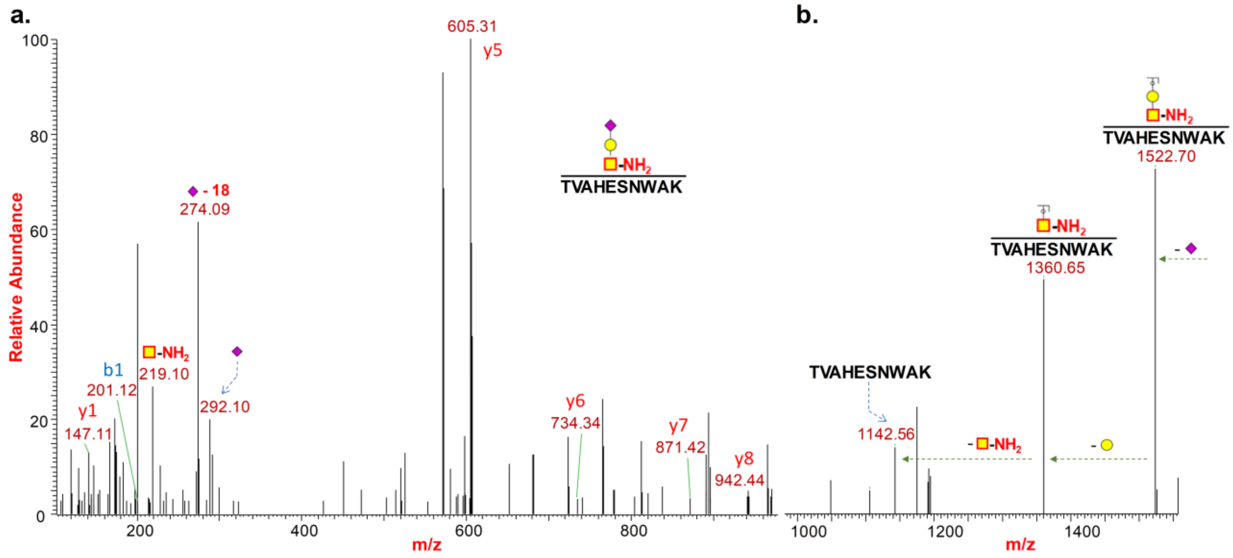


**Supp Fig S22:** HCD (a.) and CID (b.) $MS^2$ spectrum of glycopeptide TVAHESNWAK from Podocalyxin (O00592) protein from $Ac_4GalNAz$ treated PC-3 cells showing GalN-$NH_2$ incorporation on core-1 monosialylated O-glycans (On6) by the presence of oxonium ion (m/z 219.10) and neutral losses (m/z 218.09).

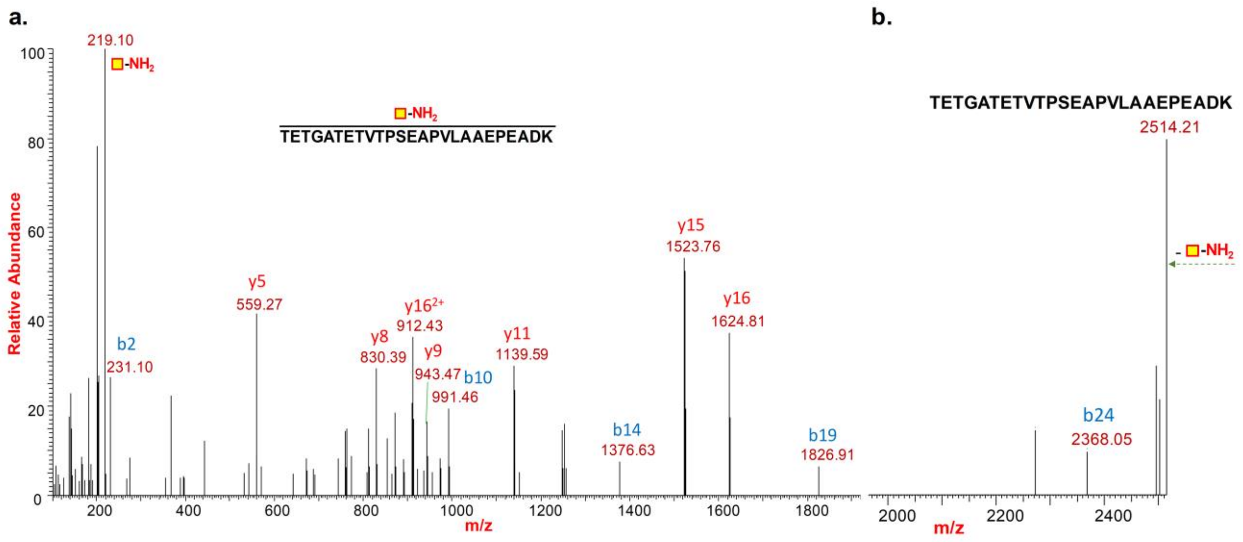


**Supp Fig S23:** HCD (a.) and CID (b.) MS2 spectrum of glycopeptide TETGATETVTPSEAPVLAAEPEADK from Thioredoxin domain-containing protein 5 (Q8NBS9) protein from Ac4GalNAz treated PC-3 cells showing GalN-$NH_2$ incorporation as Tn O-glycans (On4) by the presence of oxonium ion (m/z 219.10) and neutral losses (m/z 218.09).

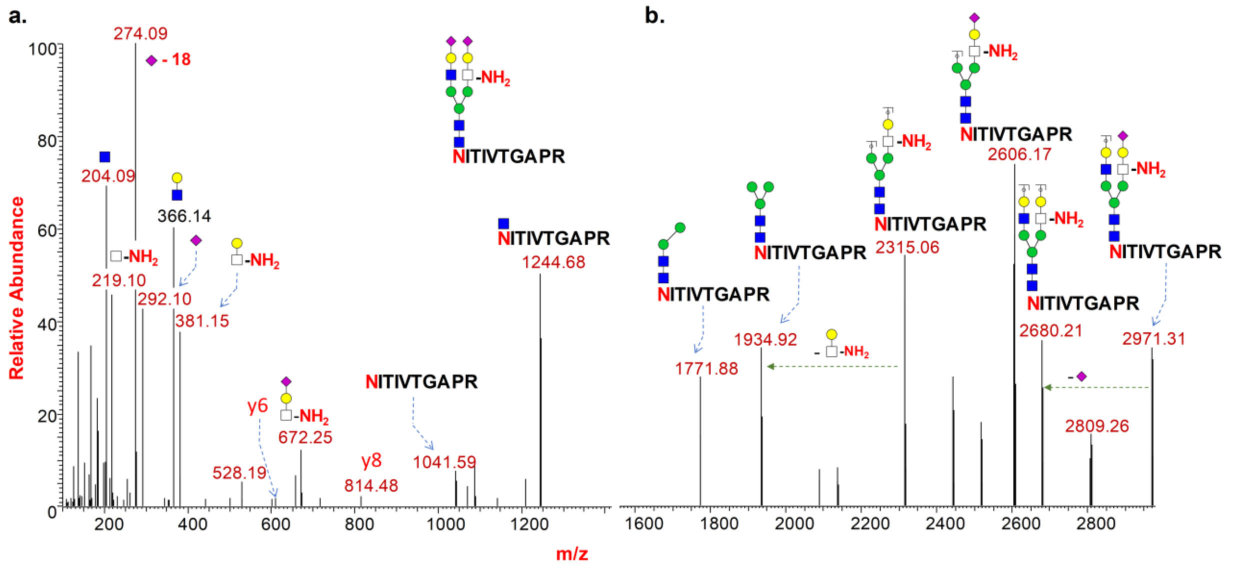


**Supp Fig S24:** HCD (a.) and CID (b.) $MS^2$ spectrum of glycopeptide NITIVTGAPR from Integrin alpha-3 (P26006) protein from $Ac_4$GalNAz treated PC-3 cells showing GlcN-$NH_2$ incorporation on the terminal of N-glycans (Nn38) and not at the core by the presence of oxonium ion (m/z 219.10, 381.15, & 672.25) and neutral losses (m/z 380.14).

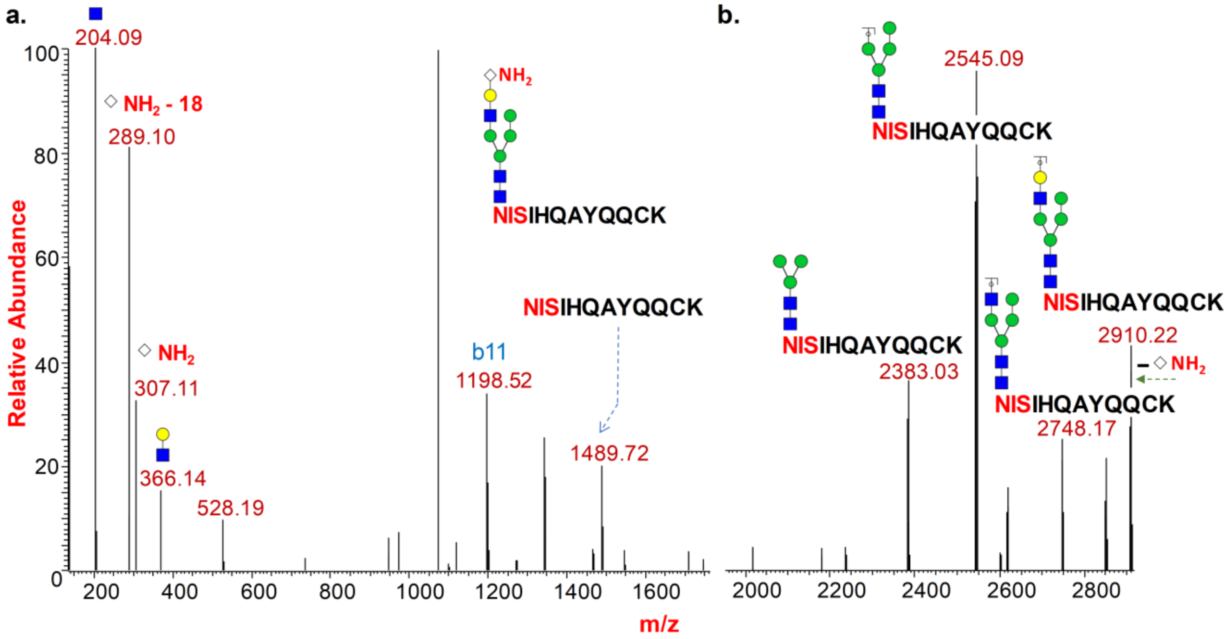


**Supp Fig S25:** HCD (a.) and CID (b.) MS[2] spectrum of glycopeptide NISIHQAYQQCK from Prominin-2 (Q8N271) protein from $Ac_4GlcNAz$ treated PC-3 cells showing $Neu5NH_2$ incorporation on the terminal of N-glycans (Nn37) and not at the core by the presence of oxonium ion (m/z 307.11 & 289.10) and neutral losses (m/z 306.10).

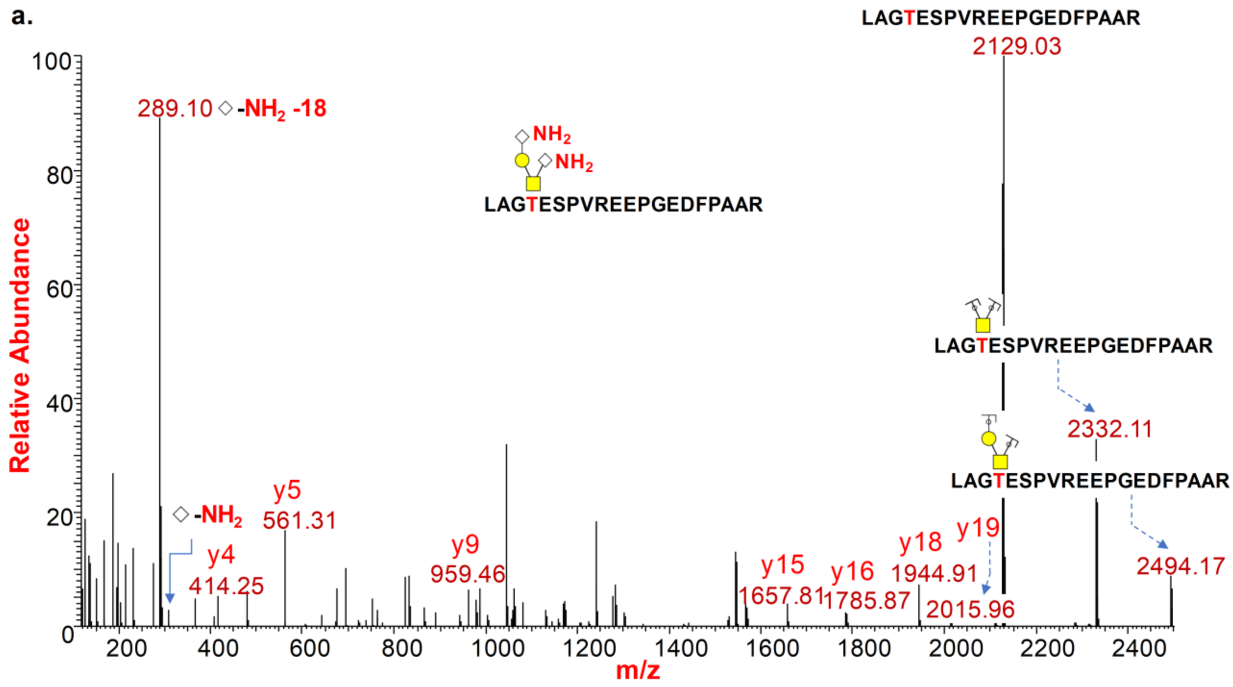

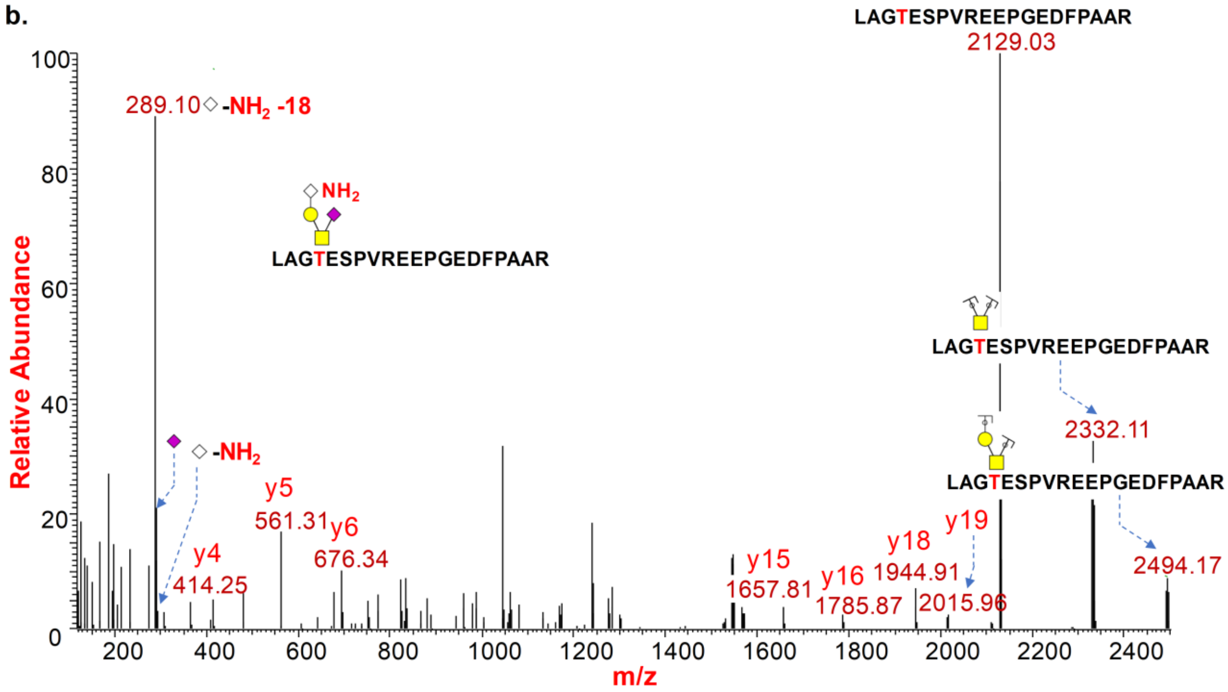


**Supp Fig S26:** HCD MS$^2$ spectrum of glycopeptide LAGTESPVREEPGEDFPAAR from Transferrin receptor protein 1 (P02786) protein from $Ac_4GlcNAz$ treated PC-3 cells **a.** showing two $Neu5NH_2$ incorporation on core-1 disialylated O-glycans (On10); **b.** showing incorporation of one $Neu5NH_2$ and one Neu5Ac on core-1 disialylated O-glycans (On9).

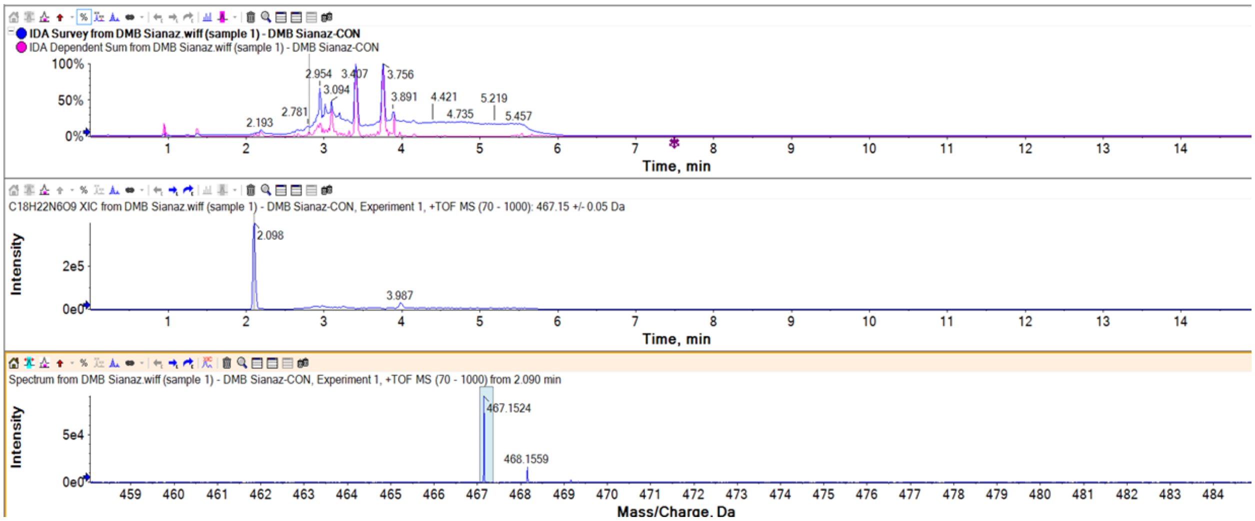


**Supp Fig S27:** Detection of DMB-SiaNAz on Jurkat cells treated with Ac4GlcNAz. The DMB-SiaNAz is confirmed by the mass spectrum and this confirms the conversion of GlcNAz to SiaNAz.

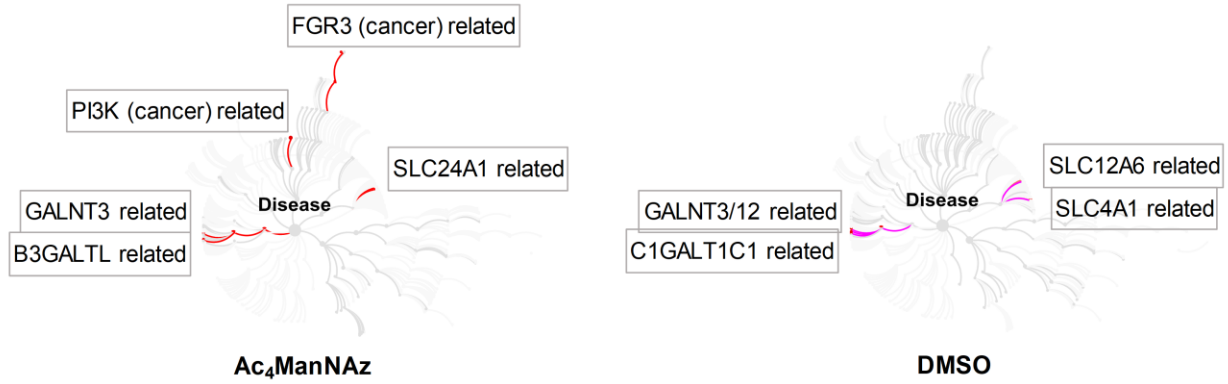


**Supp Fig S28:** Pathway analysis of the quantitative sialoglycoproteomics data from PC-3 cells in diseases revealed enrichment of Pi3K (7 proteins) and FGR3 (5 proteins) pathway only on $Ac_4ManNAz$ (total 258 proteins, Supp Table 1) and not on DMSO (total 285 proteins, Supp Table 2) treated case. (Analyzed by Reactome database with the total intensity of each detected sialoglycoprotein).[6]

### Supp References


1. Zaro, B. W., Batt, A. R., Chuh, K. N., Navarro, M. X., and Pratt, M. R. (2017) The Small Molecule 2-Azido-2-deoxy-glucose Is a Metabolic Chemical Reporter of O-GlcNAc Modifications in Mammalian Cells, Revealing an Unexpected Promiscuity of O-GlcNAc Transferase, *ACS Chem Biol 12*, 787-794.
2. Woo, C. M., and Bertozzi, C. R. (2016) Isotope Targeted Glycoproteomics (IsoTaG) to Characterize Intact, Metabolically Labeled Glycopeptides from Complex Proteomes, *Curr Protoc Chem Biol 8*, 59-82.
3. Friedman, D. B. (2007) Quantitative proteomics for two-dimensional gels using difference gel electrophoresis, *Methods Mol Biol 367*, 219-239.
4. Bond, M. R., Zhang, H., Kim, J., Yu, S. H., Yang, F., Patrie, S. M., and Kohler, J. J. (2011) Metabolism of diazirine-modified N-acetylmannosamine analogues to photo-cross-linking sialosides, *Bioconjug Chem 22*, 1811-1823.
5. Cartwright, I. L., Hutchinson, D. W., and Armstrong, V. W. (1976) The reaction between thiols and 8-azidoadenosine derivatives, *Nucleic Acids Res 3*, 2331-2339.
6. Fabregat, A., Jupe, S., Matthews, L., Sidiropoulos, K., Gillespie, M., Garapati, P., Haw, R., Jassal, B., Korninger, F., May, B., Milacic, M., Roca, C. D., Rothfels, K., Sevilla, C., Shamovsky, V., Shorser, S., Varusai, T., Viteri, G., Weiser, J., Wu, G., Stein, L., Hermjakob, H., and D'Eustachio, P. (2018) The Reactome Pathway Knowledgebase, *Nucleic Acids Res 46*, D649-D655.
